# *Pseudomonas aeruginosa* reaches collective decisions via transient segregation of quorum sensing activities across cells

**DOI:** 10.1101/2021.03.22.436499

**Authors:** Priyanikha Jayakumar, Stephen A. Thomas, Sam P. Brown, Rolf Kümmerli

## Abstract

Bacteria engage in a cell-to-cell communication process called quorum sensing (QS) to coordinate expression of cooperative exoproducts at the group level. While population-level QS-responses are well studied, we know little about commitments of single cells to QS. Here, we use flow cytometry to track the investment of *Pseudomonas aeruginosa* individuals into their intertwined Las and Rhl QS-systems. Using fluorescent reporters, we show that QS gene expression (signal synthase, receptor and exoproduct) was heterogenous and followed a gradual instead of a sharp temporal induction pattern. The simultaneous monitoring of two QS genes revealed that cells transiently segregate into low receptor (*lasR*) expressers that fully commit to QS, and high receptor expressers that delay QS commitment. Our mathematical model shows that such gene expression segregation could mechanistically be spurred by transcription factor limitation. In evolutionary terms, temporal segregation could serve as a QS-brake to allow for a bet-hedging strategy in unpredictable environments.

## Introduction

Bacterial cells typically communicate with each other via quorum sensing (QS) to coordinate collective behaviour at the group level^1–3^. One of the most widespread QS systems involves N-acyl homoserine lactone (AHL) signalling molecules that are produced by a signal synthase and subsequently accumulate both intra- and extracellularly^4^. Upon reaching a threshold concentration, typically at high population density, QS signals bind to their cognate QS receptors to form complexes that serve as transcriptional regulators. These signal-receptor dimer complexes upregulate the expression of signalling molecules in a positive feedback loop^5,6^. Subsequently, they upregulate the expression of a range of collective behaviour, including production of secreted proteases to digest food^7^, biosurfactants for group motility^8^, toxins to attack competitors^9^ and the formation of multi-cellular biofilms^10,11^. QS regulation plays a crucial role in determining the lifestyle of bacteria^12,13^, and their virulence in the context of infections^2,3,14^.

QS is extremely well studied in many taxa, including *Pseudomonas*, *Vibrio* and *Bacillus* spp.^4,15–17^. It is generally assumed that gene expression patterns and phenotypes observed at the group level are representative of what individual cells do within the population. However, this assumption conflicts with studies showing inter-individual differences in gene expression are common among clonal cells even under uniform environmental conditions^18–21^. An important question that therefore arises is whether and to what extent the coordinated QS response observed at the group level is driven by heterogeneous gene expression at the individual cell level. Several single-cell studies have reported considerable variation across cells in their QS gene expression. For example, high inter-individual variation in QS activation occurred at low cell density in *Pseudomonas putida*, which later on converged across cells at higher population density^11^. Another set of single-cell studies showed that inter-individual heterogeneity in QS gene expression and phenotypes persisted even at high population density in species such as *Vibrio harveyi*, *Pseudomonas syringae* and *Xanthomonas campestris*^22–24^. Additional heterogeneity could arise in cases where bacteria can have multiple QS systems, which are often regulatorily linked to one another through positive and negative feedbacks^25–27^. Here, heterogeneity in one QS system could propagate to a regulatorily linked second QS system, which could potentially lead to a segregation in gene expression activity, with the extreme case being that subgroups of cells specialize and communicate via different channels^28,29^.

Here, we aim to quantify patterns of single-cell gene expression heterogeneity in the multi-layer QS system of the opportunistic human pathogen *Pseudomonas aeruginosa*. This bacterium features two AHL-dependent QS systems, termed Las and Rhl^9,27^. Each system comprises its own AHL signal (Las: 3O-C12-HSL; Rhl: C4-HSL), its specific receptor, LasR and RhlR, and a set of downstream QS-controlled traits (e.g., LasB exoprotease and rhamnolipid biosurfactants synthesized by the RhlAB proteins). The two QS systems are arranged in a hierarchical signalling cascade where the Las system positively regulates the Rhl system through the LasIR transcriptional regulator dimer^9^. The two systems are often involved in regulating the expression of similar traits^30^, and together regulate the expression of almost 6% of the *P. aeruginosa* genome^31^.

In this hierarchical QS system, heterogeneity in gene expression can occur at the level of the signal, receptor and downstream genes, and the existing regulatory feedbacks could foster positive or negative correlations in gene expression between genes of the same or the interlinked QS system. We first designed chromosomally integrated single mCherry fluorescent reporter systems for six QS genes (Las system: *lasI*, *lasR*, *lasB*; and Rhl system: *rhlI*, *rhlR*, *rhlA*) to track and quantify temporal gene expression patterns and heterogeneity across single cells in clonal populations growing from low (QS-off) to high (QS-on) cell density. Next, we built a mathematical model of a simple QS regulatory circuit to identify the key elements and conditions required for heterogeneity to spur negative gene expression correlations (i.e., segregation into different expresser phenotypes) across cells. Finally, we designed double fluorescent reporters to monitor expression correlations of two QS genes in individual cells. We constructed signal-receptor (*lasI*-*lasR* and *rhlI*-*rhlR*) and receptor-downstream product (*lasR*-*lasB* and *rhlR*-*rhlA*) gene reporter pairs to quantify within-QS system gene expression correlations; and receptor (*lasR-rhlR*) and downstream product (*lasB*-*rhlA*) gene reporter pairs to compare across-QS systems gene expression correlations. This study design allows us to obtain a first comprehensive view on gene expression variation within the QS signalling cascade across cells and time within growing clonal populations.

## Results

### Las and Rhl systems are not required but are induced during growth in a nutrient rich environment

To study QS gene expression in *P. aeruginosa* PAO1, we required a medium where cells grow well and the QS network is induced. Figure 1 shows that these conditions are fulfilled in the nutrient rich Lysogeny broth (LB) medium during batch culture growth. When following gene expression of the QS signal synthase, the QS receptor, and the QS-regulated downstream genes of the Las and Rhl systems, we found all QS genes began to be expressed in the mid exponential phase and peaked in the stationary phase of culture growth (Fig. 1a-b). The gene expression kinetics measured with a plate reader further revealed that the promoter strengths differed across genes. For all QS genes, the expression increased gradually across time, and the induction pattern followed the expected temporal order from signal synthase, to receptor, to the downstream gene.

**Fig. 1.**
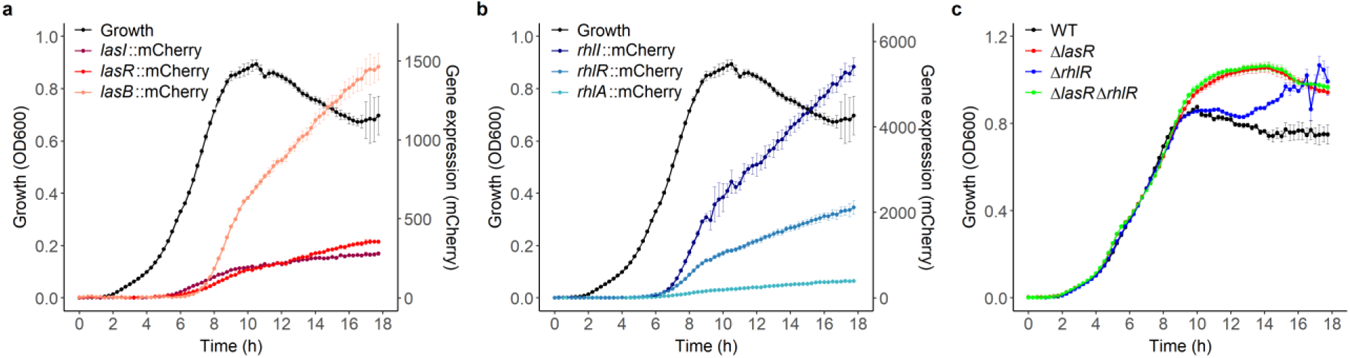
QS activity and growth in LB medium. Overlay of growth of wild type *P. aeruginosa* PAO1 strain and the expression of the QS signal synthase, the cognate QS receptor and a QS-regulated downstream gene within the Las (a) and Rhl (b) systems. Gene expression was measured as mCherry fluorescence and reported as fluorescence units, blank corrected by the background fluorescence of the wild type untagged strain. (c) Growth kinetic of QS mutants (*ΔlasR*, *ΔrhlR* and *ΔlasR-ΔrhlR* mutants) in comparison to the wild type (WT). Data shows the mean (± standard deviation) of gene expression or growth across 6 independent replicates.

We further assessed whether the QS systems are needed for growth in the nutrient rich LB environment. We used isogenic PAO1 mutants, deficient in the production of either one of the two QS receptors LasR (*ΔlasR*), RhlR (*ΔrhlR*), or both receptors (*ΔlasR-ΔrhlR*). We found that all QS mutants grew similarly compared to the wild type during the exponential phase (Fig. 1c). However, growth trajectories diverged afterwards: while the wild type strain reached stationary phase relatively early, all QS mutants continued to grow and reached significantly higher growth yields than the wild type (one-way ANOVA, F_3,20_ = 42.81, p < 0.0001, TukeyHSD pairwise comparisons: mutants vs. wild type, all comparisons P_adj_ < 0.0001). These fitness patterns suggest that expressing one or both QS systems is costly, and does not result in a net benefit during growth in LB medium. This observation is expected as LB is a rich medium and no QS-regulated traits are required for growth and survival.

### QS gene expression is switched off after constant exposure to low cell density

As the expression of QS genes peaks in the stationary phase cultures, we predicted that cells from overnight cultures are already expressing these QS genes. We indeed found this to be true for all the six QS genes in our study (Fig. 2a). However, we aimed to start our main experiments with cells that show no QS activity. To switch off the QS activity in cells, we repeatedly re-inoculated cells from overnight cultures to a low cell density environment. We observed that QS gene expression became indistinguishable from background fluorescence after two re-inoculation steps (i.e., 6 hours at low cell density, Fig. 2a), while the housekeeping genes (*rpsL, recA* and *rpoD*) remained constantly expressed (Fig. 2b). Consequently, we applied the two-step re-inoculation procedure prior to all single-cell experiments to obtain a starting population of QS-off cells.

**Fig. 2.**
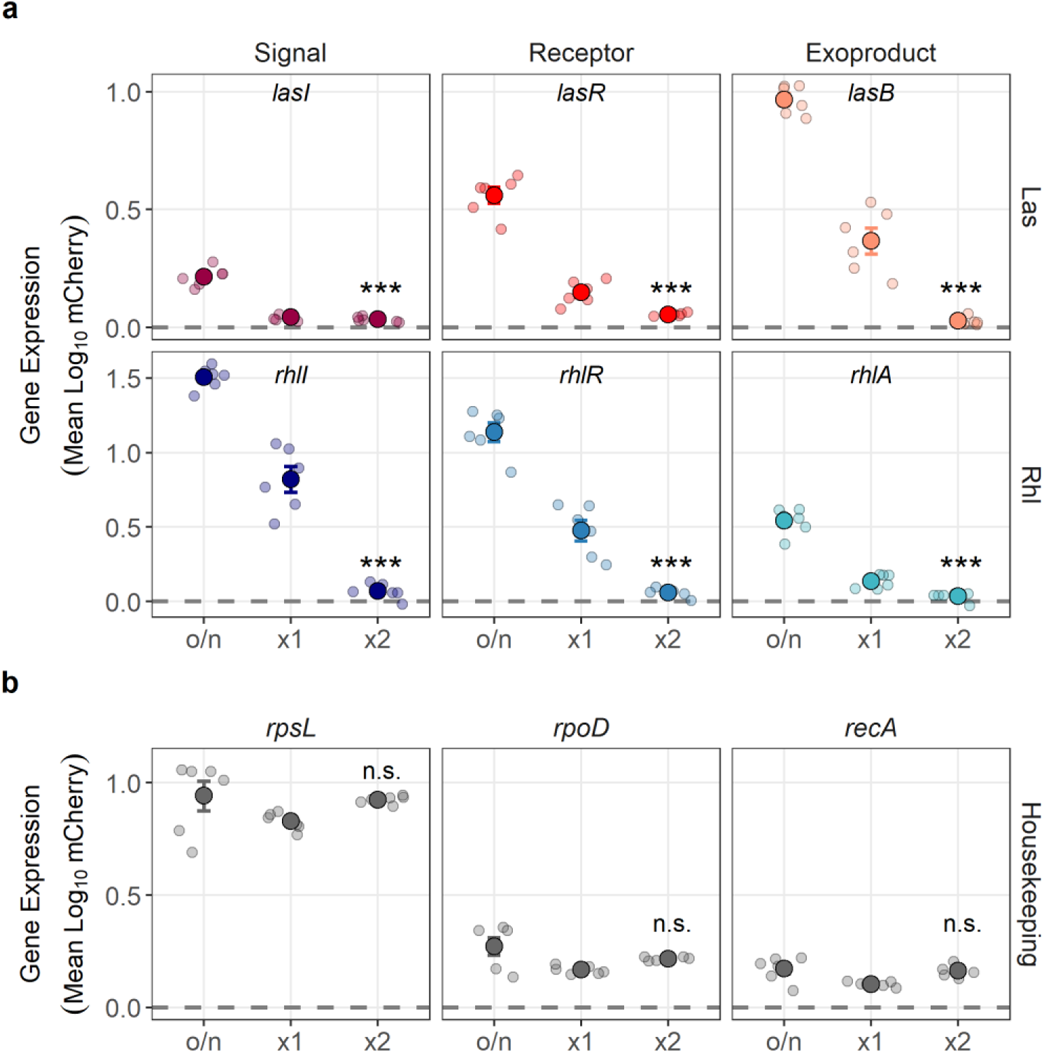
QS gene expression is switched off after prolonged exposure to low cell density. (a) Expression of QS genes in overnight cultures (o/n) and after one (x1) or two (x2) re-inoculation steps (3 hours each) in fresh LB medium. (b) Expression of housekeeping genes in cultures that underwent the same re-inoculation procedure. Dotted lines denote the mean background fluorescence. Data is the mean ± standard deviation of gene expression across 6 independent replicates (smaller circles). Asterisks indicate whether gene expression after two re-inoculations is significantly different from the expression in the overnight cultures (based on Welch’s two-sample t-tests: n.s. = not significant, *** p < 0.001).

### Single-cell expression of QS genes follows a sigmoidal increase across time

Next, we moved to the single-cell level and used flow cytometry to quantify patterns of QS and housekeeping gene expression across cells in growing clonal populations over time, starting with populations of QS-off cells at low cell density (Fig. 3). Consistent with our findings at the batch-culture level (Fig. 1), we observed a transition from an “off” (0^th^ hour) to “on” state (18^th^ hour) for the expression of all *las* and *rhl* genes. The induction of QS gene expression across time followed sigmoidal functions (Fig. 3a-b), as would be expected for a QS system. However, the transition from off to on was not sharp at a particular time point, but occurred more gradually across several hours. When looking at the distribution of gene expression across cells, we found bimodal patterns for *lasB*, *rhlI* and *rhlR*, as cells switched from off to on at different time points. In contrast, gene expression distribution was unimodal for *lasI*, *lasR* and *rhlA* at all time points (Supplementary Fig. 1a). The temporal expression patterns of QS genes differed markedly from those of the housekeeping genes, which were already expressed at 0^th^ hour and remained constitutively expressed through time (Fig. 3c).

**Fig. 3.**
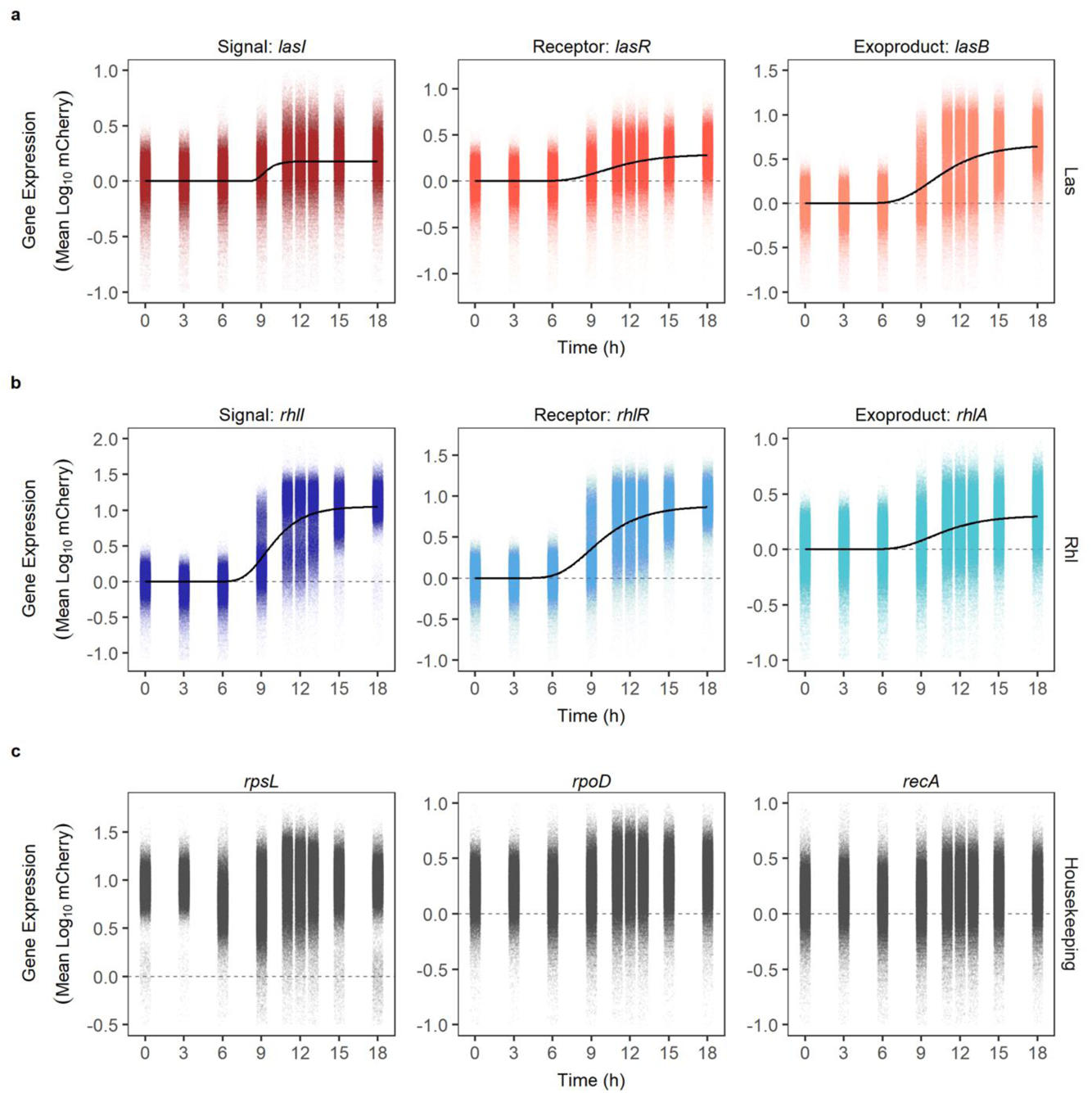
Single-cell QS gene expression patterns over time. Single-cell expression of *las* (a), *rhl* (b) and three housekeeping genes (c) over an 18-hour growth cycle in LB medium. Each dot represents a single cell (N = 50,000 per time point) detected and measured with flow cytometry. The expression of each gene was measured in wild type *P. aeruginosa* PAO1 cells with chromosomally integrated single-copy mCherry fluorescent reporters, whereby fluorescence intensity was used as a proxy for gene expression activity. Gene expression has been background subtracted by the fluorescence of the non-fluorescent wild type strain. Dashed lines at y = 0 represent the threshold value below which there is no measurable gene expression activity. Black lines show the best model fit (Gompertz function) capturing the temporal dynamics of single-cell gene expression. Because of differences in the reporter strengths, the y-axis scale varies across panels. The figure is a representative data set from one out of three complete experimental repeats (see Supplementary Fig. 2 for the other two repeats).

We then tested whether these gene expression patterns can be recovered when using GFP as the fluorescent reporter. This was possible due to our collection of double gene expression reporters (Supplementary Table 1), which had all QS genes linked to *gfp*, too. We were indeed able to recover the sigmoidal expression functions for all QS genes except *lasI* (Supplementary Fig. 2). The expression of *lasI-*GFP showed a transient increase in *lasI* expression and followed a quadratic function over time. We observed further differences between the mCherry and GFP reporters. First, fluorescence signals were picked up earlier for GFP than for mCherry, suggesting higher sensitivity for the former. Second, the bimodal gene expression patterns observed with mCherry disappeared with GFP (Supplementary Fig. 1b), indicating that the bimodality might be a specific feature of mCherry, and as a consequence of this, we refrain from providing a biological interpretation of the observed bimodality.

### QS gene expression heterogeneity peaks during the transition from exponential to stationary growth phase

We then quantified the level of heterogeneity in QS gene expression using the data from all single mCherry QS-gene reporters across the three experimental replicates. For each QS-gene reporter, time point and replicate, we calculated the coefficient of variation (CV) in gene expression across 50,000 cells (Fig. 4). Our statistical model revealed that the CV in gene expression was significantly influenced by the gene type (i.e., *las* vs. *rhl* vs. housekeeping genes; ANCOVA: F_2,234_ = 52.57, p < 0.0001), time (quadratic term: F_1,234_ = 218.34, p < 0.0001) and their interaction (F_2,234_ = 6.42, p = 0.0019). Specifically, CV was significantly higher for the *las* and *rhl* genes than for the housekeeping genes (TukeyHSD pairwise comparisons: *las* and *rhl* genes vs. housekeeping genes, both comparisons P_adj_ < 0.0001). Moreover, we observed that the CV peaked at intermediate time points (between the 9^th^ and the 11^th^ hour) for all gene groups studied, suggesting that gene expression is particularly heterogeneous during the exponential growth phase and during the activation and build-up of the QS response. CV values decreased for all genes when populations entered the stationary phase. Another way of quantifying the level of heterogeneity is to compare the standard deviations across samples, especially with log transformed data as in our case^32^. We followed this approach and recovered the same qualitative differences in single-cell gene expression heterogeneity across reporters and time points (Supplementary Fig. 4).

**Fig. 4.**
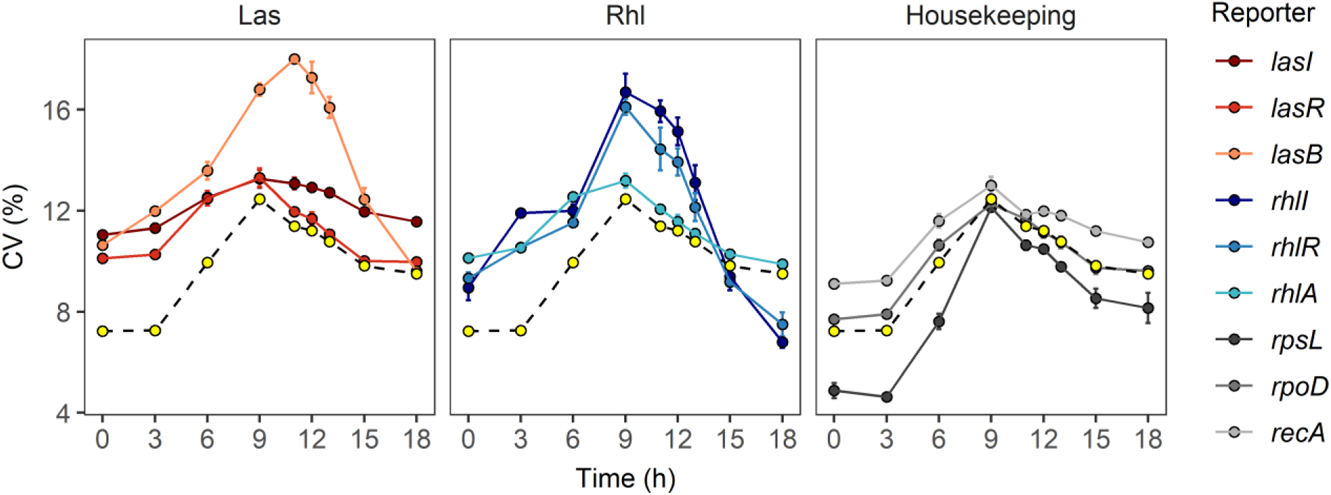
Heterogeneity in gene expression is higher for QS than for housekeeping genes and peaks at intermediate time points. Panels show the coefficient of variation (CV = percentage of the standard deviation of a sample divided by its mean) for the *las*, *rhl* and housekeeping genes. Yellow dots connected with the dashed line represent the average CV values across three housekeeping genes. Data is shown as mean ± standard error of the CV values across three independent replicates.

### Negative correlations between two QS genes can arise with limited transcription factor availability and low numbers of competing binding sites

Given the numerous positive feedback loops in the QS regulon, the null hypothesis is that the expression of any two QS genes should correlate positively across cells. The hypothesis is based on the fact that the master transcription factor (TF) of the QS regulon, the Las signal-receptor dimer, positively regulates the expression of the *lasI* signal, the *las*-downstream genes and the *rhl* regulon^3,27,33^. Accordingly, low or high TF concentration in a cell should trigger low and high expression of all QS genes, respectively, and thus lead to positive gene expression correlations across cells.

Here, we develop a mathematical model to explore whether there are conditions that could also lead to negative gene expression correlations and thus spur segregation in gene expression behaviour among cells in clonal populations. We model promoter binding and unbinding as a stochastic process, using the Gillespie algorithm^34^. Specifically, our model focuses on the Las signal-receptor dimer as the key TF in the *P. aeruginosa* QS regulon (Fig. 5a) and builds on the idea that this TF could be a limiting factor in the system^35^. In a first version of the model, we considered a simple scenario where a limited number of TFs (1 - 20) compete for the binding sites of two focal genes A and B within a cell. We consistently found that the expression of A and B correlated negatively across cells, whereby the strength of the negative association declined with increasing TF availability (Fig. 5b). This result makes intuitive sense as under stringent TF limitation, a cell can express either A or B, but rarely both genes. Next, we implemented the biologically relevant feature that in QS systems the TFs compete for more than two binding sites (up to 20 in our analysis). We observed that the strength of the negative gene expression correlations between the two focal genes declined with more competing binding sites (Fig. 5b). This result also makes intuitive sense as a limited number of TFs are distributed across many binding sites, thereby weakening gene expression correlations between any two genes within the system. Overall, our model shows that negative gene expression correlations can manifest when TF availability and the number of competing binding sites are low (Fig. 5c).

**Fig. 5.**
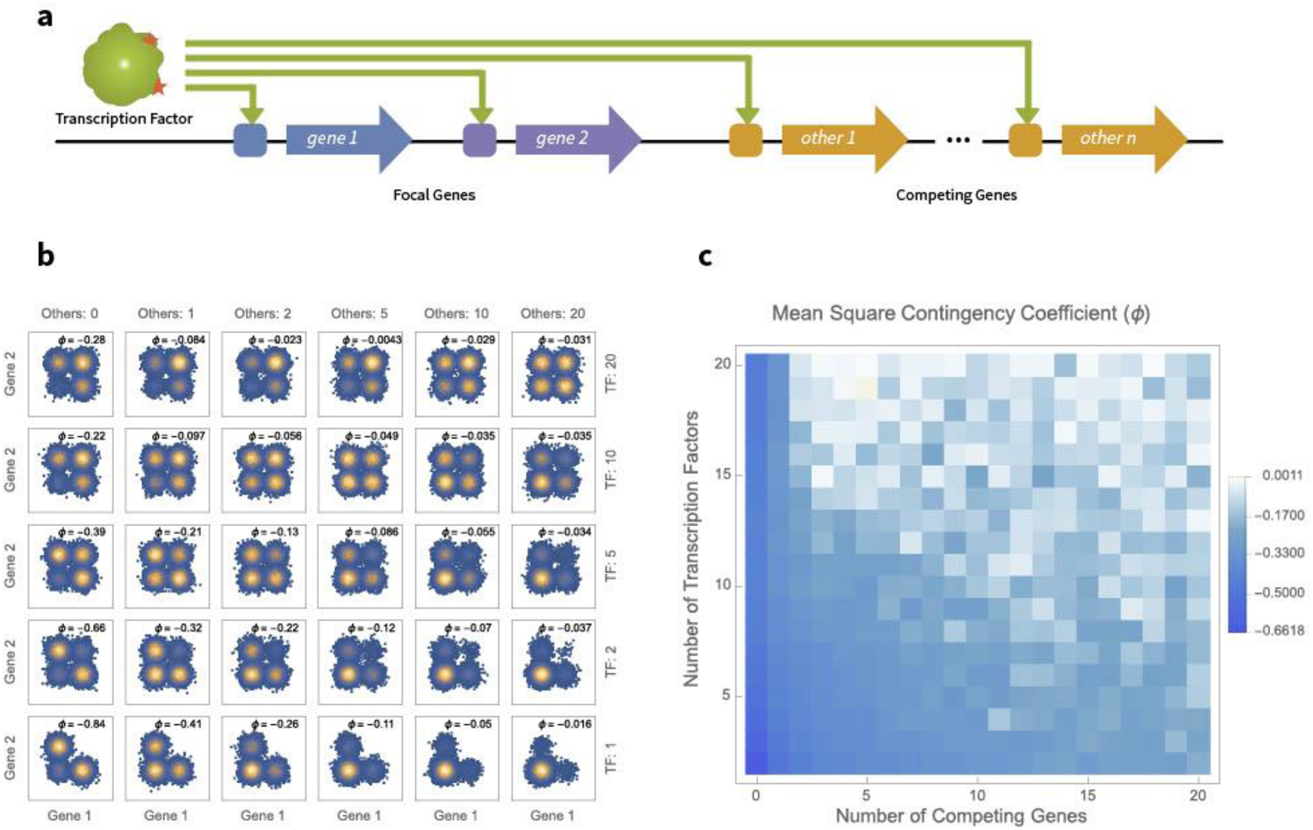
Low number of transcription factor (TF) and competing genes reveal negative correlations between the expression of two regulatorily-linked QS genes. (a) The Las signal-receptor dimer acts as a TF for two focal QS genes, as well as a number of other competing genes. (b) Relative occupancy of two focal gene promoters as a function of the number of TF complexes available and the number of additional genes competing for those complexes. Each point represents a simulated bacterial cell. Each modelled scenario is based on 20,000 simulated cells for a representative selection of parameter combinations. Gaussian noise is added to help distinguish individual cells. Correlations are quantified by the mean square contingency coefficient (ϕ), calculated without added noise. (c) Correlation heatmap between occupancy states of focal gene promoters as a function of the number of TFs available and the number of additional genes competing for the same TFs for all simulated parameter combinations.

### Characterization of the double QS gene expression reporter system

The results of the model motivated us to experimentally quantify the simultaneous investment of individual cells into two QS genes over time, and to test whether gene expression correlated positively or negatively across cells. For this purpose, we constructed a series of double gene expression reporters (Supplementary Fig. 5, Supplementary Table 1, see materials and methods), where the biologically ‘earlier’ gene in the QS cascade (e.g., *lasR*) was fused with GFP, while the biologically ‘later’ gene (e.g., *lasB*) was fused with mCherry. We conducted a series of control experiments to test the properties of our double reporter constructs. First, we confirmed that the promoterless double reporter does not show any fluorescence activity (Supplementary Fig. 5a). Second, we demonstrated that there is no fluorescence leakage between the two fluorescence channels (Supplementary Fig. 5b-c). This is important as the two reporters are sequentially arranged and activity in one promotor could trigger the expression of both fluorescence genes.

Finally, we tested for positive correlations between the GFP and mCherry signals in constructs where both fluorescent genes were fused to the same promoter. The first test involved the housekeeping gene, *rpsL*, whose promoter was fused to both GFP and mCherry. We found positive correlations between the two fluorescence signals for all time points (Fig. 6a and Supplementary Fig. 6). The Spearman’s rank correlation coefficient was highest during the early hours (ρ = 0.87) and lowest at intermediate hours (ρ = 0.55). The second test involved a QS gene, *lasB*, whose promoter was fused to both GFP and mCherry (Fig. 6b and Supplementary Fig. 6). Here, we observed no correlation between the two fluorescence readouts during the early hours (up to the 6^th^ hour, ρ = 0.03), as the gene is not expressed. At later time points, strong positive correlations built up between the two fluorescence signals (ρ = 0.69). These control experiments yielded three pieces of information. (i) The observed ρ-values were consistently below one, suggesting that there is substantial intrinsic noise in promoter activities within each cell. (ii) The GFP signal appeared earlier than the mCherry signal (Fig. 6b, 6^th^ vs. 9^th^ hour). (ii) Bimodal gene expression patterns were only observed with mCherry but not with GFP. The two latter points confirmed our observations made with the single-gene reporters.

**Fig. 6.**
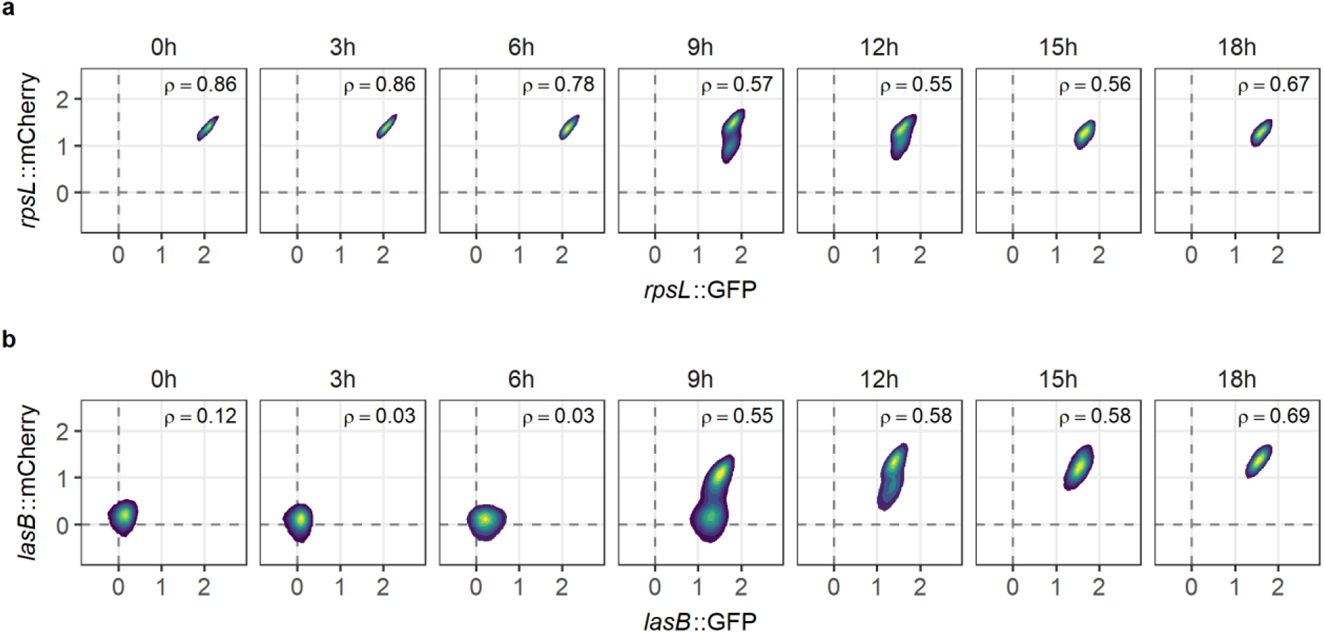
Double reporter control experiments reveal both positive correlations of gene expression and substantial intrinsic noise. Double gene expression reporters, where the promoter of the same gene – (a) the housekeeping gene, *rpsL*; (b) the QS gene, *lasB* – were fused to both the GFP and mCherry. Constructs were chromosomally integrated as single copies into the *P. aeruginosa* PAO1 wild type. (a) For *rpsL*, positive correlations were observed for all time points with an intermittent decline in the strength of the association during the exponential growth phase. (b) For *lasB*, the gene was initially not expressed, but positive correlations built up over time. Spearman correlation coefficient (ρ) below one indicates that there is substantial intrinsic noise in gene expression. Fluorescence data across 50,000 single cells are shown as 2D density plot, where yellow and blue areas represent the densest and least dense regions, respectively. Data stems from one representative experiment out of a total of three independent replicates.

### Coordinated QS response involves transient segregation in gene expression activities

We then used our main collection of double reporters to quantify the investment of individual cells into the expression of two QS genes. For all the gene pairs studied (Supplementary Table 1), we found weak or no correlations in the fluorescence signals across QS genes during the early hours of the growth cycle (Fig. 6, up to the 6^th^ hour). This is expected as the QS genes are not expressed during this period. From the 9^th^ hour onwards, QS genes began to be expressed, and a diverse set of correlation patterns ranging from no, to positive and negative correlations in gene expression across cells arose (Fig. 7).

**Fig. 7.**
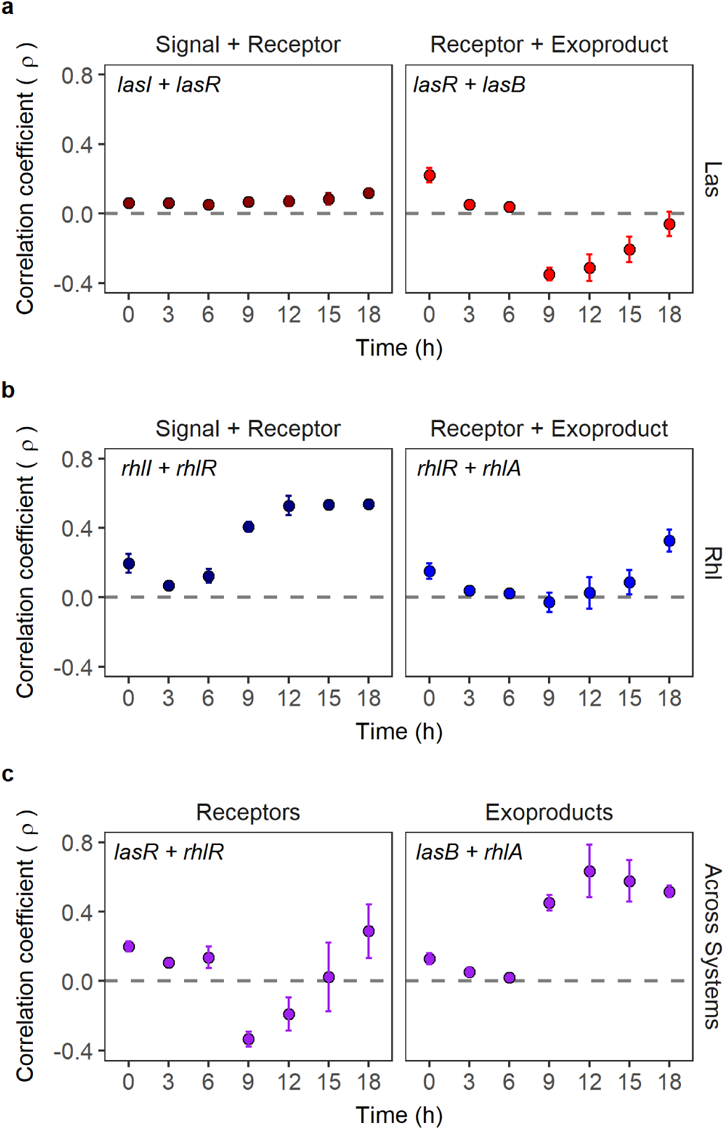
Strong positive correlations of QS downstream gene expression involve a transient segregation of cells with regard to *lasR* expression. Double gene expression reporters were used to simultaneously quantify the investment of single cells into two QS genes. Panels show the correlation in gene expression across 50,000 cells for (a) the Las-QS system: signal to receptor and receptor to downstream genes; (b) the Rhl-QS system: signal to receptor and receptor to downstream genes; (c) between QS-systems: Las to Rhl receptor and Las to Rhl downstream genes. Correlations are calculated using Spearman’s rank correlation coefficient (ρ), which can range from −1 (max. negative correlation) to 1 (max. positive correlation). Data points show means ± standard deviation across three independent replicates.

In the Las system (Fig. 7a), there were very weak positive correlations between the expression of *lasI-*GFP and *lasR-*mCherry across cells at all time points, and negative correlations in the expression of *lasR*-GFP and its downstream gene *lasB*-mCherry, especially at intermediate time points (9^th^-15^th^ hour). These patterns suggest that subpopulations of cells transiently segregate into either high *lasR* or high *lasB* expressers. In the Rhl system (Fig. 7b), there were strong positive correlations between the expression of *rhlI*-GFP and *rhlR*-mCherry from intermediate time points onwards, while a delayed build-up of positive correlations occurred in the expression of *rhlR*-GFP and *rhlA*-mCherry (Fig. 7b).

We then examined gene expression patterns across the Las and Rhl systems (Fig. 7c). We found negative correlations between the expression of *lasR*-GFP and *rhlR*-mCherry at intermediate time points (9^th^-12^th^ hour), which then became weakly positive at later time points. As above, the negative correlations indicate that cells within the clonal population transiently segregate into either high *lasR* or high *rhlR* expressers. In contrast, we found strong positive correlations between the expression of the QS downstream genes, *lasB*-GFP and *rhlA*-mCherry, a pattern we also confirmed with the marker swap control (*lasB*-mCherry and *rhlA*-GFP, Supplementary Fig. 7). Moreover, all cells expressed both *lasB* and *rhlA* from the 15^th^ hour onward (Supplementary Fig. 8). This shows that despite temporal segregation at the level of Las receptor expression, the end stage of all cells is full QS commitment. It is worth nothing that other links between the two QS systems exist, which we did not include here. For example, *lasB* expression is also responsive to the Rhl signal-receptor dimer^30^, albeit at a much lower affinity^36^. Hence, that is why we considered *lasB* as being a primarily Las-dependent trait, following previous studies^37,38^. In any case, such additional positive links are predicted to further contribute to the full QS commitment of all cells at the end stage, as observed in our experiments.

To follow up on the negative gene expression correlations involving LasR, we examined whether segregation in gene expression among cells occurred along a continuum from low to high *lasR* expression, or whether there were discrete subpopulations of low and high *lasR* expressers. Our analysis based on 2D density plots strongly support the latter scenario (Fig. 8). For both gene pairs (*lasR-lasB* and *lasR-rhlR*), we first observed a homogenous increase in *lasR* expression at the 6^th^ hour followed by a split into two subpopulations: (1) cells that further increased *lasR* expression, but neither expressed *lasB* nor *rhlR* (blue box in Fig. 8); and (2) cells that kept *lasR* expression constant, and started expressing *lasB* and *rhlR* (red box in Fig. 8). Overall, this suggests that there is a strong preference for cells to either highly express *lasR* or the Las-regulated downstream genes during the early activation of QS. Over time, the fraction of high *lasR* expressers in the population declined, leaving behind a rather uniform population of cells expressing all the three genes (*lasR*, *lasB*, *rhlR*).

**Fig. 8.**
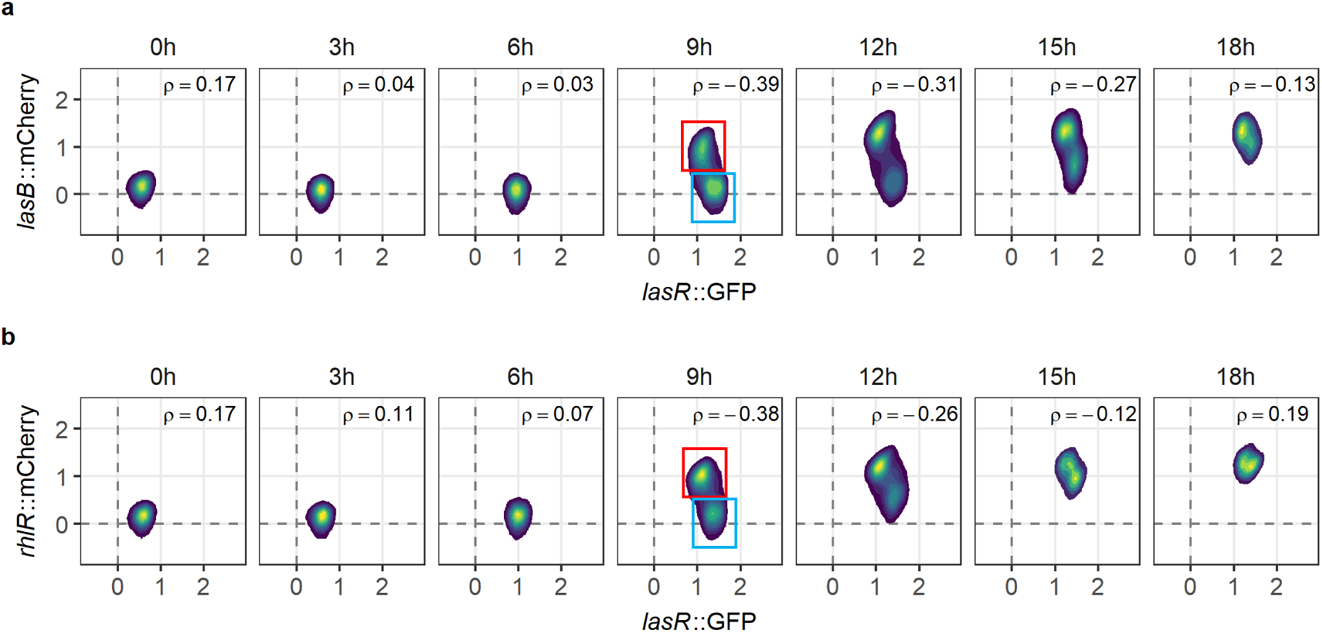
Clonal cells form subpopulations that transiently differ in their QS gene expression activities. Simultaneous single-cell expression of *lasR* and its downstream genes – (a) *lasB* and (b) *rhlR* – was measured using double fluorescent gene reporters. The population of cells began expressing *lasR* homogeneously (6^th^ hour), and split into two subpopulations, expressing either (i) high *lasR*, but no *lasB* or *rhlR* (blue box), or (ii) low *lasR* but high *lasB* and *rhlR* expression (red box). The fraction of cells belonging to subpopulation (i) declined at the later time points, resulting in one population with rather uniform gene expression activities. Fluorescence data across 50,000 single cells are shown as 2D density plot, where yellow and blue areas represent the densest and least dense regions, respectively. Dotted lines represent mean background fluorescence in the mCherry and GFP channels. Spearman’s rank correlation coefficient (ρ) between the expression of two genes are shown in each panel. Data stems from one representative experiment out of a total of three independent replicates.

## Discussion

The common view on communication via quorum sensing (QS) is that bacteria rely on the production, release and sensing of signaling molecules to trigger coordinated actions at the group level, typically at high population density^1,3,9,27,39^. As QS studies are typically conducted at batch culture levels, it is largely unknown how individual cells drive the coordinated QS responses that we observe at the group level. Here, we explored QS-behavior of individual cells of the bacterium *Pseudomonas aeruginosa*, growing from low to high cell density. We focused on the two hierarchically linked QS-systems (i.e., Las inducing Rhl)^9,27^, and indeed found high levels of coordination among cells in their final QS response. However, the way to reach this coordinated response is much more intricate than population-level observations would suggest. First, we observed that there is considerable heterogeneity among cells in their expression of Las and Rhl signal, receptor, and downstream exoproduct genes. Second, we found that gene expression heterogeneity can lead to temporal segregation in QS gene expression activities among cells. We identified the *lasR* regulator gene as the key node in the network associated with segregation. Specifically, populations temporarily split into two discrete subpopulations: low *lasR* expressers that also expressed the downstream *lasB* and *rhlR* genes; and high *lasR* expressers that expressed neither of these two downstream genes. This segregation in gene expression activity is transient and leads to a delay in the full commitment to QS at the group level. Below, we discuss potential mechanistic and evolutionary explanations for the observed transient segregation, notably whether it just reflects a by-product of the complex regulatory circuit or whether segregation could serve as an adaptive built-in brake.

At the mechanistic level, most QS systems entail multiple positive feedback loops that amplify QS sensing and response within populations of cells^40^. In the case of *P. aeruginosa*, both the Las and the Rhl systems involve such positive feedbacks, where transcription factors (i.e., signal-receptor dimers) increase the expression of their own signal synthase and their respective downstream regulons^9,41^. Thus, the null expectation would be that the expression of any two QS genes should be positively correlated across cells. We indeed observed such positive correlations for *rhl* gene pairs (*rhlI* vs. *rhlR* and *rhlR* vs. *rhlA*, Figure 7b), suggesting that this system follows the text-book QS induction model. In contrast, the Las system clearly did not follow this null model, as correlations involving the *lasR* gene were either absent (*lasI* vs. *lasR*) or negative (*lasR* vs. *lasB* and *lasR* vs. *rhlR*). Our mathematical model offers one possible solution to this conundrum by revealing that transcription factor (TF) limitation can spur negative gene expression correlations across co-regulated genes (Fig. 5). The transcription factor, Las signal-receptor dimer, is the central node in the QS-network of *P. aeruginosa.* As the supply of active TF depends on both Las signal availability and LasR copy number, it is conceivable that its supply might be particularly low early on during QS induction^35^. Moreover, TF availability might be further restricted through negative-feedback loops, such as the one operating via QslA, a repressor of LasR^27,42^. Our model further reveals that negative correlations are particularly strong when the number of genes competing for the transcription factor is low. While the number of competing genes is supposedly high for QS, the question is whether transcriptional co-factors can temporally modulate binding priorities such that the effective number of competing genes is low during QS initiation but increasing over time thereby weakening negative gene expression correlations. Altogether, our model of TF limitation offers a parsimonious explanation for the observed transient segregation among cells within the QS circuit of *P. aeruginosa*. But note that our model is also valid for other types of mechanistic constraints, such as limitations in building blocks or energy required for launching the QS cascade.

A further mechanistic aspect to consider is that segregation into subgroups can only occur when there is heterogeneity in gene expression in the first place. How does this variation come about? Our control experiments, where we fused the same promoter to two different fluorophores (mCherry or GFP) within the same cell revealed substantial intrinsic noise in gene expression (Fig. 6). This is especially true during the exponential growth phase, where gene expression correlation coefficients were moderately high (ρ = 0.56 - 0.58), given the null expectation of ρ = 1.00 for purely deterministic gene expression^18,19,43^. A second source contributing to gene expression heterogeneity could stem from extrinsic factors. While the growth conditions in our experiment were rather uniform (shaken cultures), there is still potential for stochasticity. For example, there might be stochastic inter-cell differences in nutrient uptake, which can spur variation in metabolic states across cells and induce heterogeneity in overall gene expression activity^44,45^. Moreover, the very nature of QS systems operating via diffusible signals can promote extrinsic noise. The reason is that there is likely stochasticity not only in the production but also in the uptake of signals^1,30,46^. As QS signals are at the top of the QS cascade, heterogeneity in signal availability across cells could propagate through the QS cascade, such that cells start committing to QS at different time points. Given these considerations, there seem to be ample opportunities for heterogeneity in QS gene expression to arise among cells.

From an evolutionarily perspective, it remains to be elucidated whether the observed temporal segregation in gene expression activity is an adaptive strategy, or simply a byproduct of the complex regulatory QS circuit. For example, a common pattern observed for other traits is that all cells in a population follow the same linear gene expression trajectory, but vary in the time they do so^47,48^. In our QS system, such a temporal trajectory could involve the initial onset of *lasR* expression followed by an increase in the expression of its downstream genes, *lasB* and *rhlR,* which in turn causes a drop in *lasR* expression activity. While variation in the time cells embark on this trajectory is a plausible explanation for the observed gene expression segregation, not all our data necessarily fit this scenario. For example, the gene expression trajectories observed between the 6^th^ and the 9^th^ hour suggests a bifurcated rather than a linear trajectory, where a fraction of cells (Fig. 8, red box) keeps its *lasR* expression constant and starts expressing *lasB* and *rhlR* from the 9^th^ hour onwards, while the second group (Fig. 8, blue box) increases the expression of *lasR* without launching *lasB* and *rhlR* expression. Irrespective of linear or bifurcated trajectories, the transition from the 12^th^ hour onwards, from high to low *lasR* expression and the onset of *lasB* and *rhlR* expression is slow and spurs a gradual QS induction.

One possible evolutionary advantage of the observed segregation in gene expression could be that the delay in full QS commitment serves as an built-in brake enabling clonal groups to follow a bet-hedging strategy^21,46,49^. Launching the full QS response is costly (Fig. 1c) and implies a major shift in gene expression and metabolic activities^31^. In an unpredictable environment, where conditions may rapidly change from QS-beneficial (high cell density) to non-beneficial (low cell density) situations, it could be adaptive to have a built-in brake that delays full QS-commitment of all cells in a population. In our specific case, it would mean that the high *lasR* expressers delay commitment to QS and would thus be able to quickly transit to the QS-off lifestyle should cell density diminish due to environmental changes. Fluctuating environments are common in natural settings, and bet-hedging allows cells and groups to cope and react to sudden changes in their environment^50,51^. Non-QS-related cases of bet-hedging have been demonstrated in several bacterial taxa in the context of static vs. shaken culturing conditions^52^, carbon storage and starvation^53^, and nitrogen fixation^54^. A bet-hedging strategy was also proposed as a potential explanation for the presence of QS-responsive and non-responsive subpopulations in *P. syringae* and *X. campestris*^24^. Here, we add to this body of work, by proposing that a putative bet-hedging mechanism could operate at the level of the main QS-regulator (LasR), a hypothesis that requires rigorous testing in future studies.

Finally, our study using double fluorescent gene reporters also yielded two technical insights. First, we observed that the GFP signal appears earlier than the mCherry signal for the same QS gene. This is an important factor to consider when investigating temporal succession of gene expression, as comparisons can only be made between genes linked to the same fluorophore (e.g., Fig. 1). To cope with this issue in our double fluorescent gene reporters, we fused the biologically earlier gene to GFP and the biologically later gene to mCherry. Second, we observed bimodality in the expression of certain QS genes only with the mCherry but never with the GFP marker (Supplementary Fig. 1). While this difference was not a problem for our study focusing on gene expression correlations, there is a risk of over-interpreting bimodality patterns when exclusively using single gene-reporters (i.e., mCherry in our case). Given that transient bimodal gene expression also arose for the housekeeping gene *rpsL* (Fig. 6a), we believe that bimodality could be an mCherry-specific feature. Altogether, our work with double fluorescent gene reporters strongly suggests that fluorophore combinations have to be carefully chosen to avoid misinterpretation of temporal gene expression patterns.

In conclusion, our single-cell study reveals high heterogeneity in the expression of all QS genes during the onset of QS, and transient segregation of cells in their expression levels of *lasR* versus the downstream regulated genes *lasB* and *rhlR*. Both effects together resulted in a sigmoid, yet slowly progressing QS responses at the population level. Thus, our findings support a graded^33,55^ and not an all-or-none gene expression transition^40,56,57^. While our findings support the function of QS as a means to coordinate collective actions at the group level, the path to reach collective decisions is much more intricate than population-level observations would suggest. Particularly, our insight that transient segregation delays full QS commitment raises the question of whether the delay operates as a built-in adaptive brake as part of a bet-hedging strategy or whether it is an unavoidable consequence of the complex regulatory circuit involving multiple feedback loops. While only further research can tell us more, clear is that the observed segregation leads to a gradually progressing instead of a sharp QS response at the population level.

## Materials and Methods

### Bacterial strains and reporter construction

We used *P. aeruginosa* PAO1 wild type strain (ATCC 15692) and its direct derivatives for all our experiments and *Escherichia coli* CC118 λpir for all cloning work (see Supplementary Table 1-2 for a full list of strains and plasmids used, respectively). Moreover, we used clean deletion mutants deficient in the receptor of one (PAO1 *ΔlasR*, PAO1 *ΔrhlR*) or both (PAO1 *ΔlasR ΔrhlR*) QS systems, constructed in our PAO1 background. For tracking temporal gene expression profiles, we engineered transcriptional reporter fusions in which the promoter of a gene of interest is fused to a fluorescent gene marker, *mCherry*. A single copy of the reporter construct was chromosomally integrated in the PAO1 wild type (WT) background at the neutral *att*Tn7 site using the mini-Tn7 system^58^. The detailed protocol is described elsewhere^59^. In brief, promoter regions of our genes of interest were amplified using the primers listed in Supplementary Table 3. The promoter regions include the start codon and the first ~100 base pairs of the genes, and a stop codon (TCA) was added in the reverse primers to separate the promoter regions from the *mCherry* fluorescent gene. PCR amplified promoter regions were first inserted into the pUC18-mini-Tn7-Gm-*mCherry* vector scaffold (consisting of an empty promoter site fused to *mCherry*) using restriction enzyme sites BamHI and HindIII^59^, and transformed into *E. coli*. The mini-Tn7 plasmid containing the construct was then extracted and integrated into wild type *P. aeruginosa* via electroporation. From Rezzoagli *et al.*^59^, we used the PAO1 WT::*lasR*-*mcherry* and PAO1 WT::*rhlR*-*mcherry* reporter strains. Here, we constructed four additional gene reporters (PAO1 WT::*lasI*-*mcherry,* PAO1 WT::*lasB*-*mcherry,* PAO1 WT::*rhlI*-*mcherry*, PAO1 WT::*rhlA*-*mcherry*). Moreover, we also constructed gene expression reporters for three housekeeping genes (PAO1 WT::*rpsL*-*mcherry*, PAO1 WT::*rpoD*-*mcherry*, PAO1 WT::*recA*-*mcherry*), which we used as controls for constitutively expressed genes. Sequences of our single mCherry reporter constructs were verified by DNA Sanger sequencing (Microsynth, Switzerland).

Next, we constructed a double fluorescent gene expression reporter system in the PAO1 WT background, which allows us to simultaneously measure the expression of two different genes within a single cell (Supplementary Fig. 9). The double reporter construct entails two sequential elements with identical structure. The first element is the *gfp* coding sequence (*gfpmut3*) that starts with the first promoter insertion site fused to *gfp* and ends with a terminator region. The second element is the *mCherry* coding sequence that starts with the second promoter insertion site fused to *mCherry* and ends with a terminator region. To construct the double reporter scaffold, we used the pUC18-mini-Tn7-Gm-mCherry vector scaffold from the single reporter construct (from above), which already has a promoter insertion site (with restriction enzymes BamHI and HindIII) fused to *mCherry*. The terminator region was inserted downstream of *mCherry* using restriction enzyme sites NsiI and SacI. The *gfp* sequence was amplified from pEX-A128-GFPmut3 using primers with two unique restriction enzymes each (listed in Supplementary Table 3) and inserted upstream of our *mCherry* construct with the restriction enzymes sites HindIII and KpnI using a T4 Ligase enzyme (Thermo Fischer Scientific). The introduction of two additional restriction enzyme sites, SpeI and PacI in these primers allows the insertion of a promoter region upstream of *gfp* (using restriction enzymes KpnI and PacI) and insertion of terminator region downstream of *gfp* (using restriction enzymes HindIII and SpeI). Promoter regions of genes of interest (*lasI, lasR, lasB, rhlI, rhlR* and *rhlA*) were amplified using primer pairs listed in Supplementary Table 3 with restriction enzyme sites KpnI and PacI (for *gfp* fusion), or BamHI and HindIII (for *mCherry* fusion). The promoter regions are the same as described above for the single *mCherry* reporter constructs. Ribosomal binding sites (RBSs) were added upstream of the start codons of *gfp* and *mCherry.* The terminator region consists of four rho-independent terminators^18^ and was inserted (i) in between the *gfp* and *mCherry* constructs (to avoid cross fluorescence of *gfp* into the *mCherry* channel) and (ii) downstream of the *mCherry* construct (to minimise the differences in the *gfp* and *mCherry* transcriptional complexes). After the insertion of promoter regions of genes of interest, the double reporter constructs were cloned in the mini-Tn7 system in *E. coli* and later integrated into *P. aeruginosa* wild type cells via electroporation. A detailed step-by-step cloning protocol is described elsewhere^59^. Sequences of our double reporter constructs were verified by DNA Sanger sequencing (Microsynth, Switzerland).

As controls for leaky expression in our double fluorescent gene reporter construct, we constructed: (1) promoterless *gfp* and *mCherry* strain (PAO1 WT::*empty-gfp-empty-mCherry*); (2) constitutively-expressing *gfp* but promoterless *mCherry* strain (PAO1 WT::*rpsL-gfp-empty-mCherry*); and (3) promoterless *gfp* but constitutively-expressing *mCherry* (PAO1 WT::*empty-gfp-rpsL-mCherry*). For our study design, we constructed six QS double reporters to measure simultaneous expression of: (1) signal and receptor within Las and Rhl systems (PAO1 WT::*lasI-gfp-lasR-mCherry;* PAO1 WT::*rhlI-gfp-rhlR-mCherry*); (2) receptor and downstream product within Las and Rhl systems (PAO1 WT::*lasR-gfp-lasB-mCherry;* PAO1 WT::*rhlR-gfp-rhlA-mCherry*); (3) receptors across Las and Rhl systems (PAO1 WT::*lasR-gfp-rhlR-mCherry*); and (4) downstream products across Las and Rhl systems (PAO1 WT::*lasB-gfp-rhlA-mCherry*). The order of QS genes in our double reporter constructs was determined by the biological order of these genes within the QS cascade: from signal to receptor and subsequently to its downstream product within QS systems; and from Las to Rhl across systems. Because the GFP protein matures faster than the mCherry protein, we always fused the biologically earlier gene (e.g., *lasR*) within a given QS gene pair to *gfp*, and the biologically later gene (e.g., *lasB*) to *mCherry.* To explore the properties of our double reporter construct, we engineered three control strains: (1) PAO1 WT::*rpsL-gfp-rpsL-mCherry*, for which we expect strong positive correlations of the fluorescent signals across all time points, and deviation from a perfect correlation would provide a measure of intrinsic noise; (2) PAO1 WT::*lasB-gfp-lasB-mCherry*, for which we expect strong positive correlations of the fluorescent signals to build up over time when the QS cascade is induced; (3) PAO1 WT::*rhlA-gfp-lasB-mCherry*, a promoter swap control for which we expect the same type and strength of correlation as for PAO1 WT::*lasB-gfp-rhlA-mCherry*. We regard (3) as an ideal control as both genes encode downstream products of the Las and Rhl systems, and we can thus verify that there is no marker effect at the end point of the QS cascade.

### Growth conditions

Overnight cultures were inoculated from single bacterial colonies and grown in 6mL Lysogeny broth (LB), at 37°C, 220 rpm for 18 hours. Prior to experiments, overnight cultures were washed twice with 0.8% NaCl and adjusted to an optical density (OD, measured at 600 nm) of 1.0. All growth and gene expression assays were performed in LB medium. For this, cells from overnight cultures were inoculated into fresh LB medium to a final starting OD_600_ of 0.01. For population-level experiments, cells were grown in 96-well plates containing 200 µL of LB medium per well and incubated at 37°C. For single-cell experiments, cells were grown in 24-well plates containing 1.5 mL of LB medium per well and incubated at 37°C and 170rpm. For all cloning work, we used ampicillin (100 µg/µL) to select for *E. coli* transformants carrying the mini-Tn7 plasmid containing the fluorescent gene reporter constructs, and gentamicin (30 µg/µL) to select for *P. aeruginosa* transformants containing the integration of fluorescent gene reporters. LB and antibiotics were purchased from Sigma-Aldrich, Switzerland.

### Population level growth and gene expression assays

We first conducted a population level experiment to verify that QS genes are expressed in *P. aeruginosa* in LB medium. For this, we used all six single QS gene mCherry reporters and the untagged PAO1 wild type strain (without mCherry reporter). We inoculated cells from overnight cultures into fresh LB medium to a final starting OD_600_ of 0.01 in individual wells on 96-well plates as described above. Plates were incubated at 37°C in a multimode plate reader (Tecan Infinite M-200, Switzerland). We measured mCherry fluorescence (excitation: 582 nm, emission: 620 nm) and growth (OD_600_) every 15 minutes (after a shaking event of 15 seconds) over a duration of 18 hours. Subsequently, to remove background fluorescence, we calculated the mean mCherry fluorescence intensity per time point of the untagged PAO1 wild type strain and subtracted these values from the measured mCherry fluorescence values of the QS gene reporter strains across time points. In a second experiment, we tracked the growth of PAO1 wild type strain and the three QS mutants to assess the fitness consequences of being QS-deficient in our experimental medium. Cells were prepared and grown as described above. With the plate reader, we measured growth (OD_600_) every 15 minutes (after a shaking event of 15 seconds) for a total of 18 hours. Total growth yield for each strain was calculated as a measure of fitness.

### Repeated re-inoculation of cultures to switch off QS

To ensure that QS is switched off prior to the start of our single-cell experiments, we re-inoculated cells into fresh LB medium twice to consistently keep them at low cell density. This is necessary because cells in overnight bacterial cultures are in the late stationary phase, and thus most likely express QS genes. Therefore, we took the six single QS gene reporters and the three housekeeping gene reporters and did the following. We inoculated cells from overnight cultures into fresh LB medium to a final starting OD_600_ of 0.01 in individual wells of 24-well plates as described above. Plates were incubated at 37 °C and 170 rpm. After 3 hours, 100 µL of bacterial cell cultures were transferred into 1.5 mL of fresh LB medium per well on a new 24-well plate. 100 µL of these diluted bacterial cell cultures were removed for gene expression measurement with the flow cytometer (see detailed protocol below). Plates were incubated in the same condition as above for another 3 hours. Then, we again transferred 100 µL of bacterial cell cultures into 1.5 mL of fresh LB medium per well on a new 24-well plate, and 100 µL of these diluted bacterial cell cultures were removed for gene expression measurement with the flow cytometer. The remaining bacterial cell cultures (i.e., cells in 1.5 mL of LB medium per well) were the starting population (t = 0h) for all our single cell experiments. As this protocol proved useful in switching off QS gene expression, we applied it to all subsequent single-cell gene expression experiments.

### Time-resolved QS gene expression at the single cell level

Next, we followed single-cell gene expression of QS genes using all the six single mCherry fluorescent gene reporters starting at t = 0h (where the QS systems are off) grown in 1.5 mL of LB medium in 24-well plates, incubated at 37 °C and 170 rpm over the course of 18 hours. We also measured gene expression of the housekeeping gene reporters. Due to our experimental design requiring time-dependent gene expression measurements, we split our single-cell experiments into two time blocks (i.e., on two separate days). Specifically, we set up a first set of plates with cell cultures at t = 0h, incubated the plates as described above, and measured gene expression at time points 0^th^, 3^rd^, 6^th^ and 9^th^ hour on day 1. Subsequently, we set up a second set of plates with cell cultures at t = 0h, incubated them as described above, and measured gene expression at time points 11^th^, 12^th^, 13^th^, 15^th^ and 18^th^ hour on day 2. In the follow up experiments, we used the QS double fluorescent gene reporters, along with the controls described above, to track simultaneous single-cell expression of two QS genes within and across Las and Rhl systems. The cell cultures were set up as above, except that we reduced the time points at which we measured gene expression (i.e., 0^th^, 3^rd^, 6^th^, 9^th^ on day 1; and 12^th^, 15^th^ and 18^th^ on day 2). Gene expression was measured with the flow cytometer (see detailed protocol below). For this, at time point t = 3h, we removed an aliquot of 100 µL of the growing cell cultures to measure gene expression in undiluted samples. At time point t = 6h, an aliquot of 50 µL of cell cultures was removed, and at time point t = 9h and beyond, an aliquot of 10 µL of cell cultures was removed, and cells were diluted in appropriate amount of 1X filter sterilized phosphate buffer saline (PBS, Gibco, ThermoFisher, Switzerland) for gene expression measurement. The above-described procedure was repeated three independent times.

### Flow cytometry to measure single-cell fluorescence gene expression

We used FACSymphony cell analyzer (BD Bioscience, Flow Cytometry Facility, University of Zurich) to measure single-cell GFP (blue laser, excitation at 488 nm, 530/30 filter) and mCherry fluorescence (yellow-green laser, excitation at 561 nm, 610/20 filter). We recorded 50,000 events per sample (for each strain and for each time point) with a low flow rate. The threshold of particle detection was set to 200 V for the forward and side scatter in the Cytometer Setup and Tracking (CS&T) settings of the instrument. In the first set of experiments using single mCherry reporter strains, we used PAO1 WT::*empty-mCherry* as a non-fluorescent control (as a measure of background fluorescence) and the three single constitutively expressed housekeeping gene reporters as positive control for mCherry fluorescence. In the subsequent experiments with double fluorescent gene reporters, we used the PAO1 WT::*empty-gfp-empty-mCherry* (non-fluorescent control as a measure of background fluorescence), PAO1 WT::*rpsL-gfp-rpsL-mCherry* (positive control for both GFP and mCherry fluorescence), PAO1 WT::*rpsL-gfp-empty-mCherry* (positive control for GFP fluorescence, negative control for mCherry fluorescence) and PAO1 WT::*empty-gfp-rpsL-mCherry* (negative control for GFP fluorescence, positive control for mCherry fluorescence).

### Flow cytometry data processing

We used FlowJo software (BD, Bioscience) for flow cytometry data analysis. To distinguish cells from background events, we gated for the most homogeneous population by applying a gate in the forward (FSC) and side scatter (SSC), and all analyses were performed on this gated population. We exported the GFP and mCherry fluorescence values for every single event recorded from FlowJo and imported into R (version 3.6.1) for further analyses and plotting. For all dataset, we first log transformed the mCherry and GFP expression values. Then, we subtracted the log transformed fluorescence values of each reporter gene at every time point by the mean of log transformed fluorescence values of the wild type non-fluorescent strains (which represents a measure of background fluorescence) for the respective time points. To be able to fit sigmoid logistic models to single-cell gene expression data, we scaled the above fluorescence values of each QS reporter gene per time point to the mean of these fluorescence values of the respective QS reporter genes at time point t = 0h. This ensures that the mean QS gene expression is zero at time point t = 0h, a key requirement for model fitting.

### Modelling

To analyze how different conditions and parameters might lead to negative correlations between gene expression levels, we developed a simple agent-based model in Mathematica version 12.1.1. The model relies on the Gillespie algorithm to simulate three coupled reactions^34^. Agents participating in the reactions include a generalized transcription factor (TF), promoters for two focal genes (pG1 and pG2), and a promoter for other competing genes (pGC). Each gene has its own reaction equation in the form of TF + pG ⇋ TFpG, with equal binding and unbinding rates. Model parameters are (1) the availability of the TF, and (2) the number of competing genes. We also varied binding and unbinding rates within and between the equations, but such variations did not change the qualitative results. The model implementation simulates 20,000 individual bacterial cells. It allows chemical reactions to proceed for a fixed time period and the code then records the reaction states at the end of this time. We considered a gene to be active if a TF is bound to its promoter and inactive otherwise. As the simulation only allows binary values (on or off) for each gene, we used the mean square contingency coefficient (ɸ) as a measure of correlation.

### Statistical analysis

We performed all statistical analyses with R studio (version 3.6.1). To compare growth of wild type and QS mutants in batch cultures, we fitted a parametric growth model (SSlogis) using grofit package in R, extracted the total growth yield of each strain and used one-way ANOVA and post-hoc Tukey’s HSD to statistically compare them. To test whether QS gene expression is switched off after re-inoculation, we used unpaired two-sample t-tests.

We fitted a series of models to estimate single-cell gene expression trajectories for each QS reporter across time. Specifically, we compared general linear models (linear and quadratic functions) with non-linear sigmoid models (gompertz function), and used the Akaike information criterion (AIC) to identify the best model fit (i.e., reflected by the lowest AIC value). This step was performed on both the mCherry fluorescence data obtained from the single mCherry reporters and the GFP fluorescence data obtained from the double fluorescent gene reporters.

In order to assess single-cell heterogeneity in QS gene expression, we calculated the coefficient of variation (CV) and standard deviation (SD) of the log-transformed fluorescence values (without background subtraction) of the six QS gene reporters for every time point. To test whether the CV and SD varies in response to the gene type (i.e., *las* genes, *rhl* genes and housekeeping genes) and time, we first performed an analysis of co-variance (ANCOVA), with gene type as a factor, and time as covariate. We used Tukey’s HSD for post-hoc pairwise comparisons between gene type. To assess the direction and strength of correlation across single cells in their expression of two genes, we calculated the Spearman’s rank correlation coefficient between the expression of GFP-tagged and mCherry-tagged genes. For all data sets, we checked whether the residuals were normally distributed by referring to Q-Q plots and by consulting the results of the Shapiro-Wilk test for normality.

## Acknowledgements

We thank Chiara Rezzaogli, Michael Weigert and Subham Mridha for help in the strain construction, Tobias Wechsler, Jos Kramer, Nils Eling and Aleix Boquet-Pujadas for help with statistical analysis and curve fitting, and the Flow Cytometry Facility of University of Zurich for technical support. This work was funded by the European Research Council (ERC) under the European Union’s Horizon 2020 research and innovation programme (grant agreement no. 681295).

## Author contributions

P.J. and R.K. designed the study, P.J. performed the experiments, S.A.T. and S.P.B. developed and analysed the mathematical model, P.J. and R.K. analysed the experimental data, and all authors interpreted the results and wrote the paper.

