## Supplemental Figures and Tables for "*Pseudomonas aeruginosa* reaches collective decisions via transient segregation of quorum sensing activities across cells"

**This file contains:**

Supplementary Figures 1 – 9

Supplementary Tables 1 – 3

Supplementary Reference

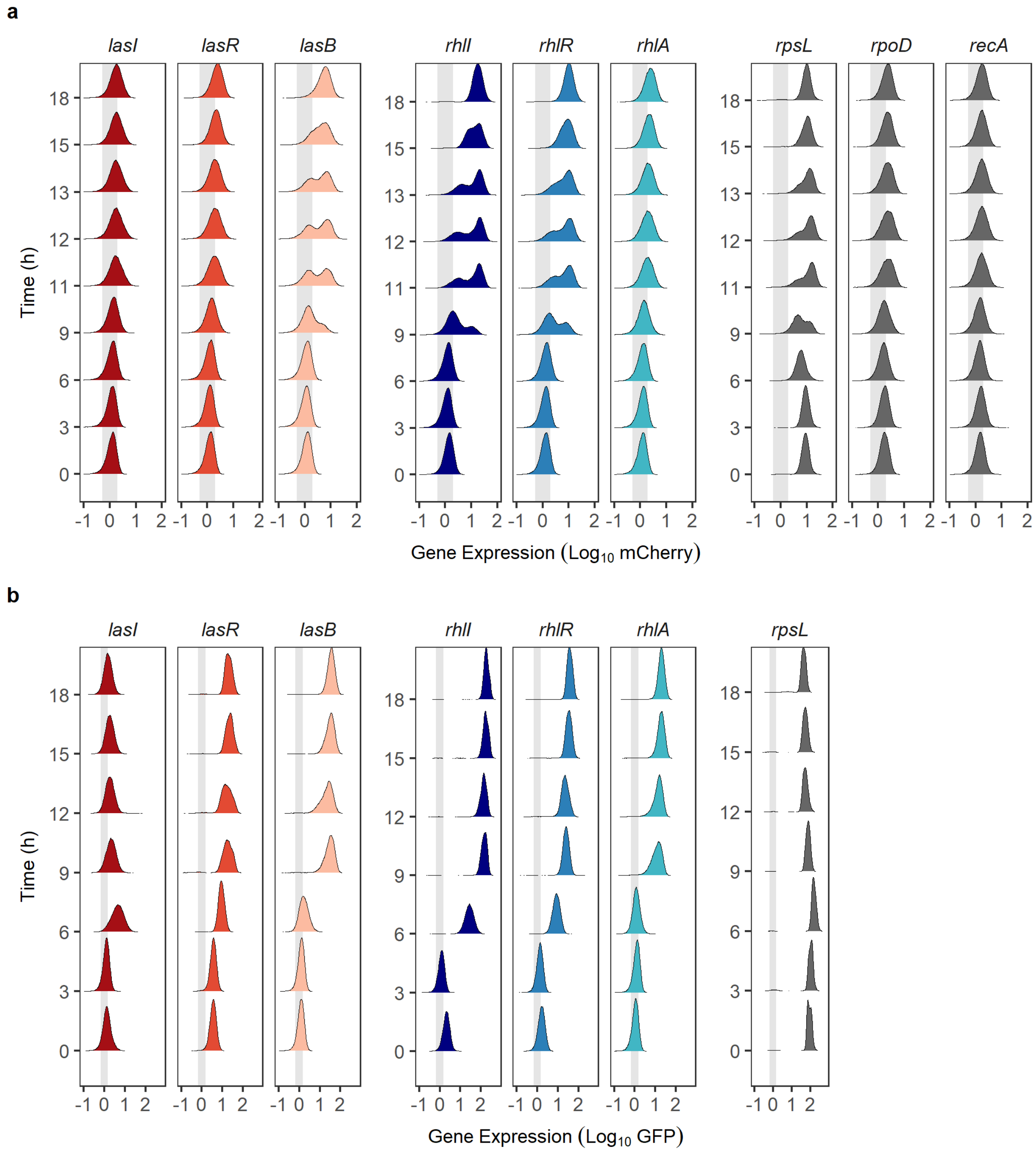


**Supplementary Figure 1.** Distribution of QS gene expression across single cells. Gene expression distribution of *las, rhl* and housekeeping genes when fused to mCherry (a) or GFP (b) in wild type PAO1 over an 18-hour growth cycle in LB medium. Distribution represents 150,000 cells per time point. Data stems from three independent replicates (i.e. Fig. 3 and Supplementary Fig. 2). Grey shaded area represents the standard deviation of background fluorescence.

**a**

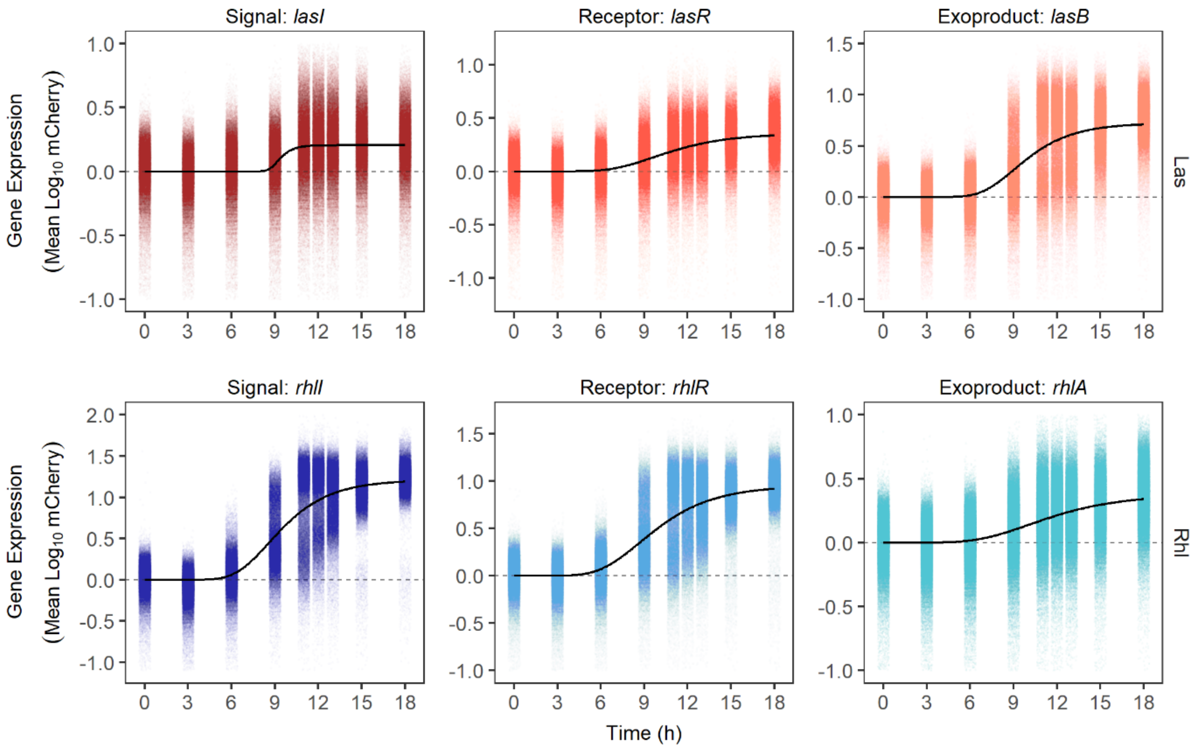
Signal: *lasI*
Receptor: *lasR*
Exoproduct: *lasB*
Signal: *rhlI*
Receptor: *rhlR*
Exoproduct: *rhlA*
Gene Expression
(Mean Log10 mCherry)
Las
Rhl
Time (h)

**b**

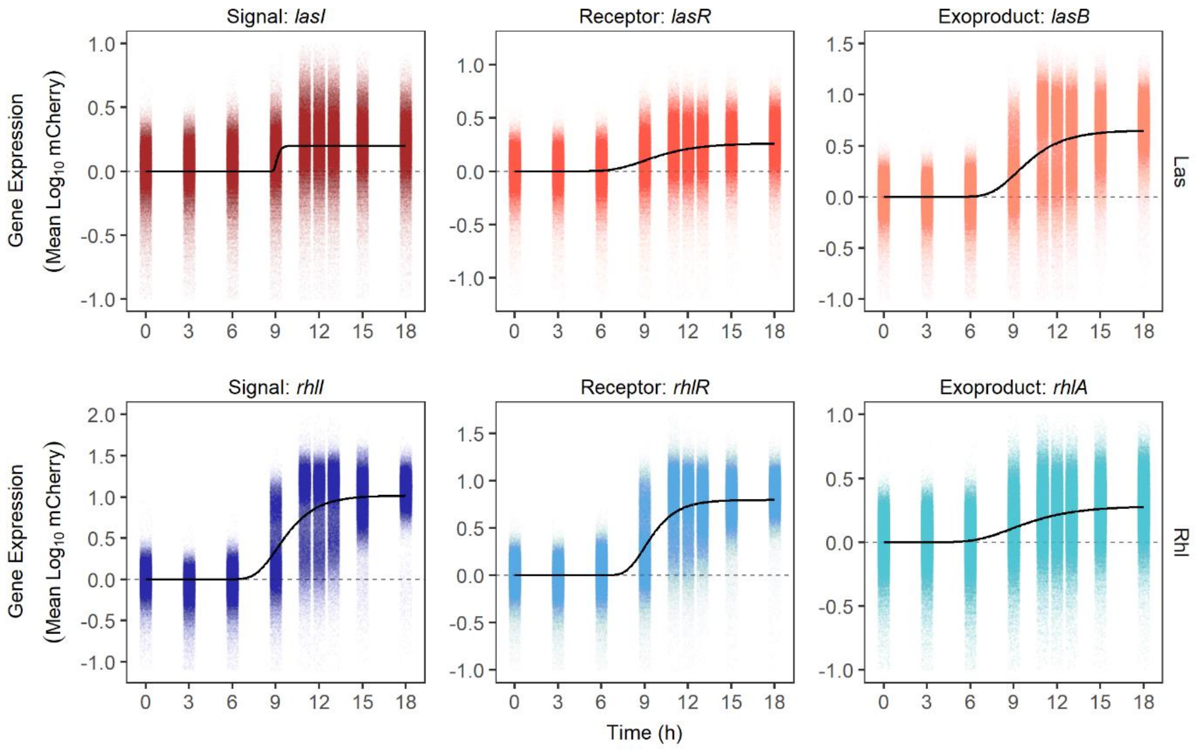
Signal: *lasI*
Receptor: *lasR*
Exoproduct: *lasB*
Signal: *rhlI*
Receptor: *rhlR*
Exoproduct: *rhlA*
Gene Expression
(Mean Log10 mCherry)
Las
Rhl
Time (h)

**Supplementary Figure 2.** Repeats of single-cell QS gene expression patterns over time. Single-cell expression trajectories of *las* and *rhl* genes for repeat 2 (a) and repeat 3 (b). See Fig. 3 for Repeat 1. Each dot represents a single cell (N = 50,000 per time point) detected and measured with flow cytometry. Dashed line at y = 0 represent the threshold value below which there is no measurable gene expression activity. Black lines show the best model fit (Gompertz function) capturing the temporal dynamics of single-cell gene expression.

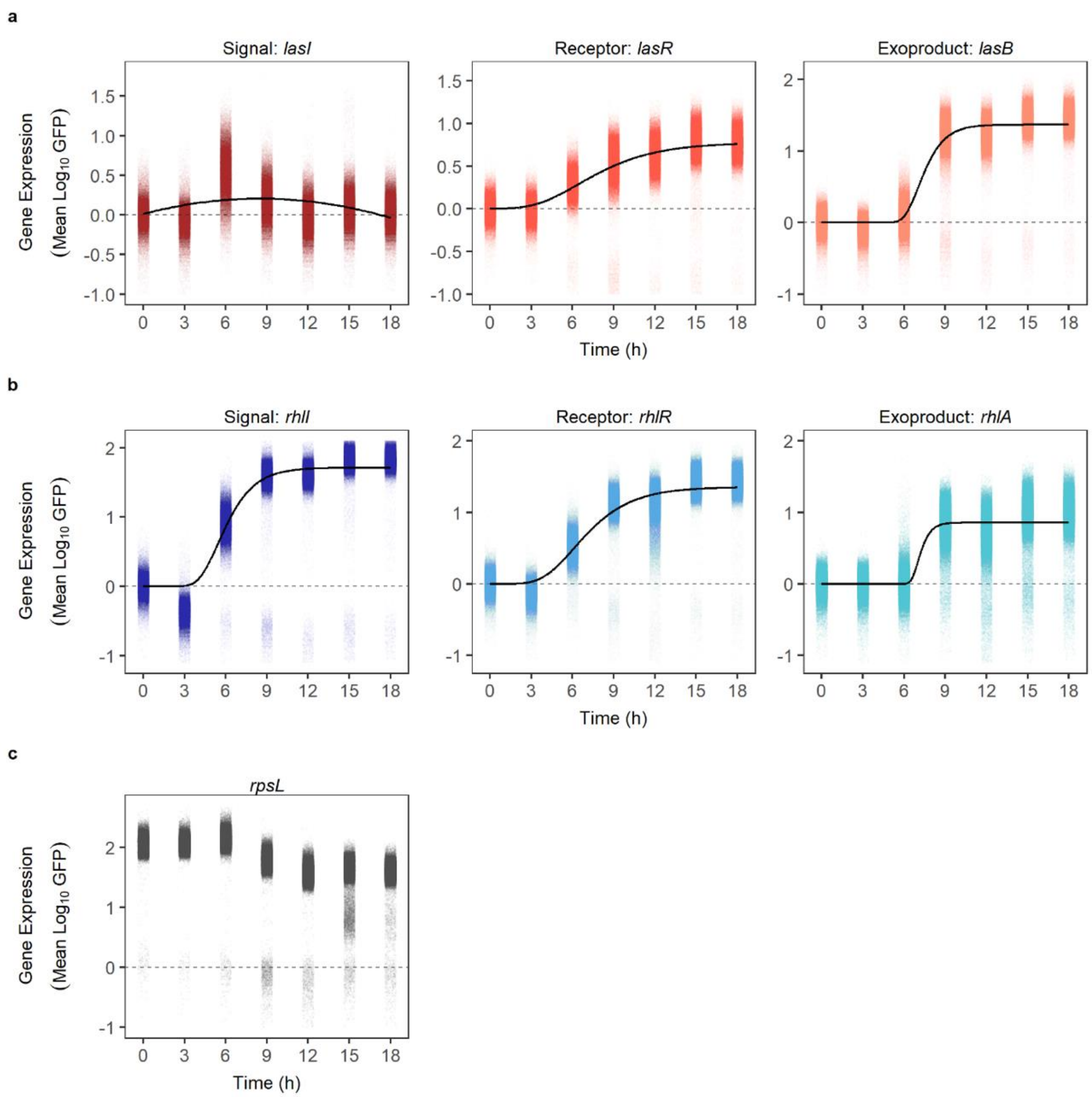


**Supplementary Figure 3.** Single-cell expression of GFP-tagged QS genes. Single-cell expression of *las* (a), *rhl* (b) and housekeeping gene, *rpsL* (c) in wild type PAO1 cells when fused with GFP in the chromosomally integrated double fluorescent gene reporters (in comparison to single mCherry reporters in Figure 3). Each dot represents a single cell (N = 50,000 per time point) detected and measured with flow cytometry. Gene expression has been background subtracted by the fluorescence of the non-fluorescent wild type strain. Dashed line at y = 0 represent the threshold value below which there is no measurable gene expression activity. Black lines show the best model fit capturing the temporal dynamics of single-cell gene expression (Gompertz function for all QS genes, except *lasI* for which a quadratic function explained most of the variation). Because of differences in the reporter strengths, the y-axis scale varies across panels. The figure shows representative data from one out of three complete experimental repeats.

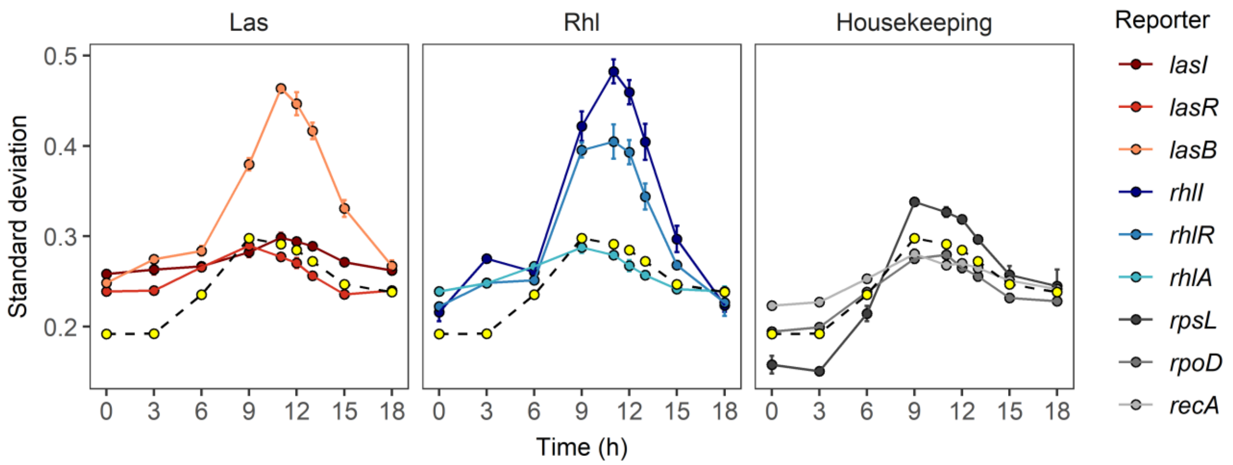


**Supplementary Figure 4.** Comparing standard deviations (SD) across cells shows that QS gene expression is more heterogenous than housekeeping gene expression. Panels show the SD of las, *rhl* and housekeeping gene expression across 50,000 cells. SD in gene expression was significantly influenced by the gene type (i.e. *las* vs. *rhl* vs. housekeeping genes; ANCOVA: $F_{2,234} = 23.48$, $p < 0.0001$), time (quadratic term: $F_{1,234} = 25.60$, $p < 0.0001$) and their interaction ($F_{2,234} = 0.71$, $p = 0.0045$). SD was significantly higher for the *las* and *rhl* genes than for the housekeeping genes (TukeyHSD pairwise comparisons: *las* and *rhl* genes vs. housekeeping genes, both comparisons $P_{adj} < 0.0001$).Yellow dots connected with the dashed line represent the average standard deviation across the three housekeeping genes. Data are shown as the mean ± standard error of the standard deviation values across three independent replicates.

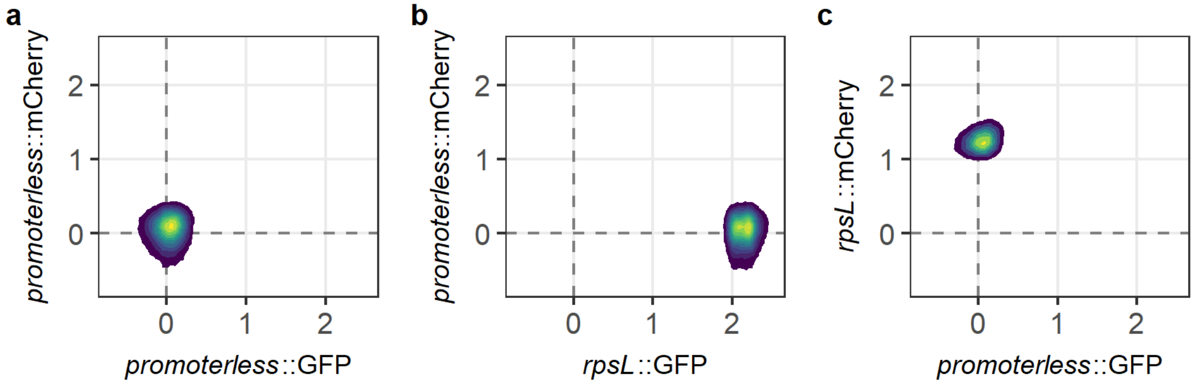


**Supplementary Figure 5.** Background fluorescence controls. Background fluorescence in the GFP and mCherry channels when measured using (a) a non-fluorescent strain carrying an integrated double reporter construct of promoterless *gfp* and *mCherry* strain (PAO1 WT::*empty-gfp-empty-mCherry*); (b) constitutively-expressing *gfp* but promoterless *mCherry* strain (PAO1 WT::*rpsL-gfp-empty-mCherry*), and (c) promoterless *gfp* but constitutively-expressing *mCherry* (PAO1 WT::*empty-gfp-rpsL-mCherry*). There was no evidence of cross fluorescence, demonstrating that the built in terminator sites ensure that the GFP and mCherry constructs are independent. Data across single cells is plotted as 2D density plot, where yellow represents the densest region and blue as the least dense region. Dotted lines represent mean background fluorescence in the mCherry and GFP channels.

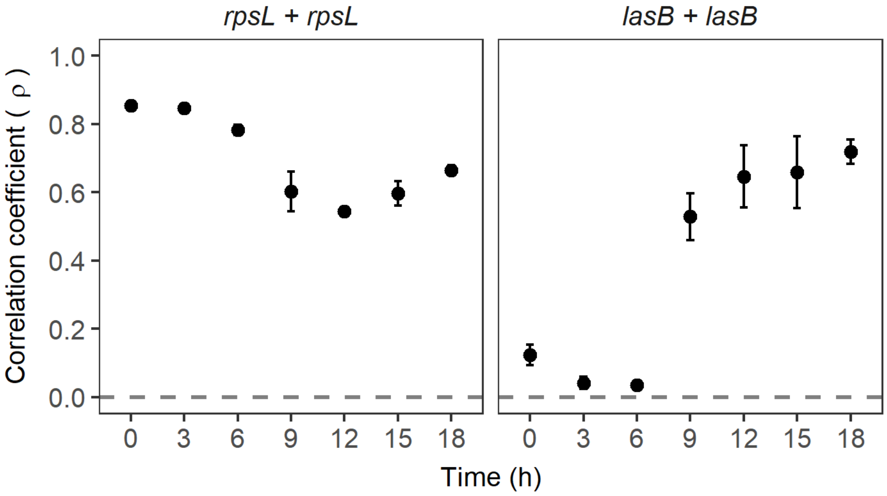


**Supplementary Figure 6.** Double reporter control experiments. Double gene expression reporters, where the promoter of the same gene – the housekeeping gene *rpsL*; and the QS gene *lasB* – were fused to both the GFP and mCherry. Panels show the averaged Spearman's rank correlation coefficient across three independent replicates (data of a single replicate is presented as a 2D density plot in Fig. 6a). Correlations are calculated using Spearman's rank correlation coefficient. Dotted lines represent zero correlation in gene expression. Data show the mean ± standard deviation of the correlation coefficient across three independent replicates.

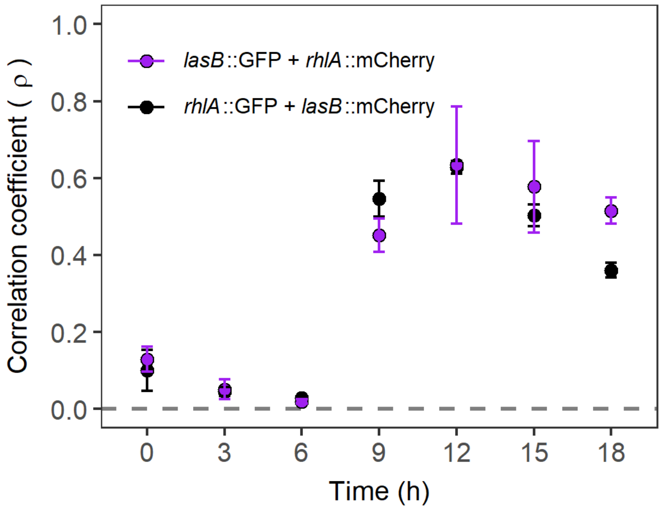


**Supplementary Figure 7.** Swapped fluorescent gene reporter control. Correlation coefficient of QS gene expression when fluorescent gene reporters are swapped in the double reporter construct (black). Correlations are calculated using Spearman's rank correlation coefficient. Dotted lines represent zero correlation in gene expression. Data shows the mean ± standard deviation of correlation coefficient across three independent repeats.

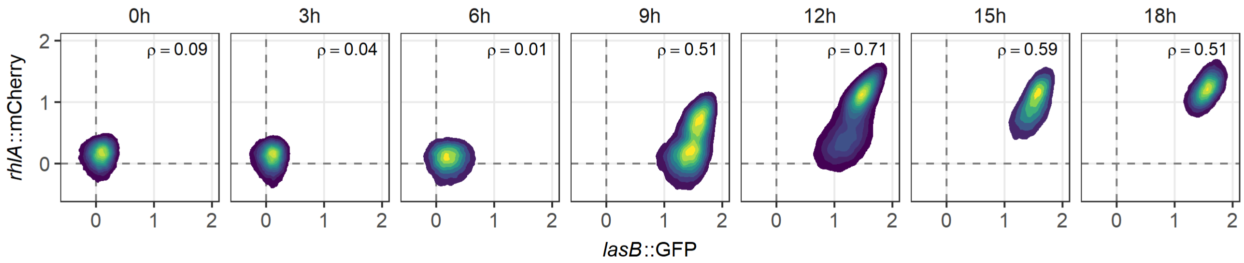


**Supplementary Figure 8.** End level Las- and Rhl-QS commitment is positively correlated across cells. Simultaneous single-cell expression of Las- and Rhl-regulated traits (*lasB* and *rhlA*, respectively) was measured using double fluorescent gene reporters. Both genes were initially not expressed, but positive correlations built up over time as cells began to commit to QS. Fluorescence data across 50,000 single cells are shown as 2D density plot, where yellow and blue areas represent the densest and least dense regions, respectively. Dotted lines represent mean background fluorescence in the mCherry and GFP channels. Spearman's rank correlation coefficient (ρ) between the expression of two genes are shown in each panel. Data stems from one representative experiment out of a total of three independent replicates.

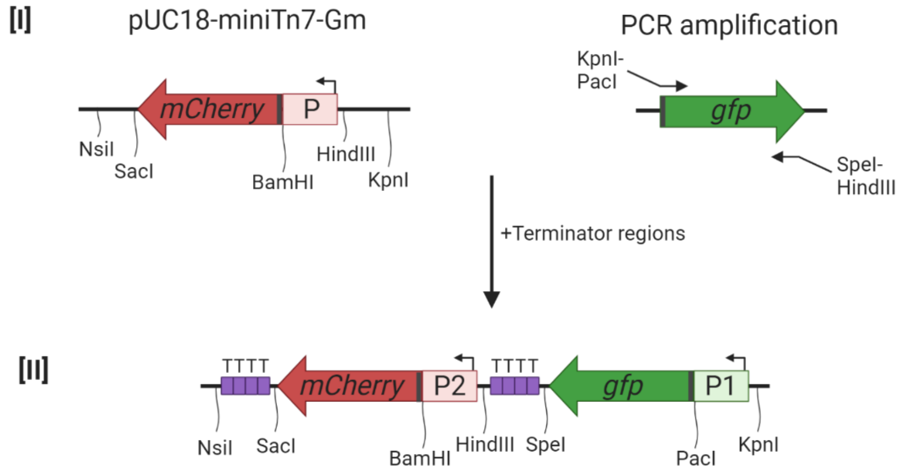


**Supplementary Figure 9.** Double fluorescent gene reporter scaffold. [I] Fluorescent gene marker *gfp* (*gfpmut3*) was PCR amplified (using primers denoted as arrows with unique restriction enzymes) and ligated to the pUC18-mini-Tn7-Gm-mCherry vector plasmid containing an empty promoter site fused to *mCherry*. [II] Promoter sites are denoted as P1 (fused to *gfp*) and P2 (fused to *mCherry*). Promoter regions of genes of interest were added at site P1 or P2 using restriction enzyme sites, KpnI and PacI, or, HindIII and BamHI, respectively. Each terminator region has four rho-independent terminators denoted as "T" (purple boxes). Ribosomal binding sites (RBSs) are shown as dark grey rectangles at the start of the fluorescent gene markers *gfp* and *mCherry*.

**Supplementary Table 1.** List of bacterial strains

| Strains | Description | Source or reference |
|---|---|---|
| ***E. coli*** | | |
| CC118 λpir | Δ (*ara, leu*)$_{7697}$ *araD*139 *ΔlacX74 galE galK phoA20 thi-1 rpsE rpoB* ($Rf^R$) *argE(am) recA1 λpir*$^+$ | [1] |
| ***P. aeruginosa*** | | |
| PAO1 wild type | Wild type strain (ATCC 15692) | This laboratory |
| PAO1 *ΔlasR* | Deficient in the receptor of Las system | This laboratory |
| PAO1 *ΔrhlR* | Deficient in the receptor of Rhl system | This laboratory |
| PAO1 *ΔlasR ΔrhlR* | Deficient in the receptor of both Las and Rhl systems | This laboratory |
| ***P. aeruginosa* wild type PAO1 (single fluorescent gene reporters)** | | |
| *promoterless-mCherry* | Non-fluorescent strain with an empty promoter site fused to mCherry | This study |
| *lasI*-mCherry | Transcriptional fusion *lasI*-mCherry from pSR01 | This study |
| *lasR*-mCherry | Transcriptional fusion *lasR*-mCherry from pSR02 | [2] |
| *lasB*-mCherry | Transcriptional fusion *lasB*-mCherry from pSR03 | This study |
| *rhlI*-mCherry | Transcriptional fusion *rhlI*-mCherry from pSR04 | This study |
| *rhlR*-mCherry | Transcriptional fusion *rhlR*-mCherry from pSR05 | [2] |
| *rhlA*-mCherry | Transcriptional fusion *rhlA*-mCherry from pSR06 | This study |
| *rpsL*-mCherry | Transcriptional fusion *rpsL*-mCherry from pSR07 | This study |
| *rpoD*-mCherry | Transcriptional fusion *rpoD*-mCherry from pSR08 | This study |
| *recA*-mCherry | Transcriptional fusion *recA*-mCherry from pSR09 | This study |
| ***P. aeruginosa* wild type PAO1 (double fluorescent gene reporters)** | | |
| *empty-mCherry-empty-GFP* | Non-fluorescent strain with an empty promoter site 1 fused to GFP, and empty promoter site 2 fused to mCherry | This study |
| *lasI-GFP-lasR-mCherry* | Transcriptional fusion *lasI-GFP* and *lasR-mCherry* from pDR01 | This study |
| *lasR-GFP-lasB-mCherry* | Transcriptional fusion *lasR-GFP* and *lasB-mCherry* from pDR02 | This study |
| *rhlI-GFP-rhlR-mCherry* | Transcriptional fusion *rhlI-GFP* and *rhlR-mCherry* from pDR03 | This study |
| *rhlR-GFP-rhlA-mCherry* | Transcriptional fusion *rhlR-GFP* and *rhlA-mCherry* from pDR04 | This study |

| | | |
|---|---|---|
| *lasR-GFP-rhlR-mCherry* | Transcriptional fusion *lasR-GFP* and *rhlR-mCherry* from pDR05 | This study |
| *lasB-GFP-rhlA-mCherry* | Transcriptional fusion *lasB-GFP* and *rhlA-mCherry* from pDR06 | This study |
| *rhlA-GFP-lasB-mCherry* | Transcriptional fusion *rhlA-GFP* and *lasB-mCherry* from pDR07 | This study |
| *rpsL-GFP-rpsL-mCherry* | Transcriptional fusion *rpsL- GFP* and *rpsL-mCherry* from pDR08 | This study |
| *lasB-GFP-lasB-mCherry* | Transcriptional fusion *lasB- GFP* and *lasB-mCherry* from pDR09 | This study |
| *rpsL-GFP-empty-mCherry* | Transcriptional fusion *rpsL-GFP* and *empty-mCherry* from pDR11 | This study |
| *empty-GFP-rpsL-mCherry* | Transcriptional fusion *empty-GFP* and *rpsL-mCherry* from pDR12 | This study |

**Supplementary Table 2.** List of plasmids

| Plasmid name | Description | Source or reference |
|---|---|---|
| pUX-BF13 | Helper plasmid to provide Tn7 transposase proteins | [3] |
| pUC18-mini-Tn7-Gm | $Gm^r$ on mini-Tn7; for chromosomal insertion in $Gm^s$ bacteria in the *att*Tn7 site | [4] |
| pEX-A128-terminators | Commercial plasmid with terminator sequences between HindIII and SpeI sites. | [5] |
| pEX-A128-GFPmut3 | Commercial plasmid with *gfp* sequence | This laboratory |
| **Single fluorescent gene reporter plasmids** | | |
| pUC18-mini-Tn7-Gm-mCherry | Derived from pUC18-mini-Tn7-Gm; $Gm^r$ on mini-Tn7; used for chromosomal insertion of *mCherry* at the *att*Tn7 site in $Gm^s$ bacteria | This laboratory |
| pSR01 | pUC18-mini-Tn7-Gm-mCherry with *lasI-mCherry* | This study |
| pSR02 | pUC18-mini-Tn7-Gm-mCherry with *lasR-mCherry* | [2] |
| pSR03 | pUC18-mini-Tn7-Gm-mCherry with *lasB-mCherry* | This study |
| pSR04 | pUC18-mini-Tn7-Gm-mCherry with *rhlI-mCherry* | This study |
| pSR05 | pUC18-mini-Tn7-Gm-mCherry with *rhlR-mCherry* | [2] |
| pSR06 | pUC18-mini-Tn7-Gm-mCherry with *rhlA-mCherry* | This study |
| pSR07 | pUC18-mini-Tn7-Gm-mCherry with *rpsL-mCherry* | This study |
| pSR08 | pUC18-mini-Tn7-Gm-mCherry with *rpoD-mCherry* | This study |
| pSR09 | pUC18-mini-Tn7-Gm-mCherry with *recA-mCherry* | This study |
| **Double fluorescent gene reporter plasmids** | | |
| pUC18-mini-Tn7-Gm-mCherry-GFP | Derived from pUC18-mini-Tn7-Gm-mCherry; with amplified GFP from pEX-A128-GFP | This study |
| pDR01 | pUC18-mini-Tn7-Gm with *lasI-GFP* and *lasR-mCherry* | This study |
| pDR02 | pUC18-mini-Tn7-Gm with *lasR-GFP* and *lasB-mCherry* | This study |

| | | |
|---|---|---|
| pDR03 | pUC18-mini-Tn7-Gm with *rhlI-GFP* and *rhlR-mCherry* | This study |
| pDR04 | pUC18-mini-Tn7-Gm with *rhlR-GFP* and *rhlA-mCherry* | This study |
| pDR05 | pUC18-mini-Tn7-Gm with *lasR-GFP* and *rhlR-mCherry* | This study |
| pDR06 | pUC18-mini-Tn7-Gm with *lasB-GFP* and *rhlA-mCherry* | This study |
| pDR07 | pUC18-mini-Tn7-Gm with *rhlA-GFP* and *lasB-mCherry* | This study |
| pDR08 | pUC18-mini-Tn7-Gm with *rpsL-GFP* and *rpsL-mCherry* | This study |
| pDR09 | pUC18-mini-Tn7-Gm with *lasB-GFP* and *lasB-mCherry* | This study |
| pDR10 | pUC18-mini-Tn7-Gm with *rpsL-GFP* and *lasI-mCherry* | This study |
| pDR11 | pUC18-mini-Tn7-Gm with *rpsL-GFP* and *empty-mCherry* | This study |
| pDR12 | pUC18-mini-Tn7-Gm with *empty-GFP* and *rpsL-mCherry* | This study |

**Supplementary Table 3.** List of primers. Restriction enzyme sites are underlined in the sequences.

| Primer | Sequence (5'-3') | Template DNA | Construct |
|---|---|---|---|
| **For cloning promoter region upstream of mCherry** | | | |
| lasI_Fwd_HindIII | CAGTAAGCTTGCCCGGAAGGCC ATGTTTTG | PAO1 gDNA | pSR01 |
| lasI_Rev_BamHI | CAGTGGATCCTCATGGCCAGT GGTATCGAGAAT | PAO1 gDNA | pSR01 |
| lasR_Fwd_HindIII | ATGACAAGCTTTGGAAAAGTGG CTATGTCGC | PAO1 gDNA | pDR01 |
| lasR_ Rev_BamHI | CGTTCGGATCCTTAGGCGCTCC ACTCCAATTTTC | PAO1 gDNA | pDR01 |
| lasB_Fwd_HindIII | CAGTAAGCTTCGAATTCCGCGG CCAGGAAAGCGTGCAA | PAO1 gDNA | pSR03, pDR02, pDR07 and pDR09 |
| lasB_Rev_BamHI | CAGTGGATCCTCAATCGAACAAC AGGTCAAGCGTA | PAO1 gDNA | pSR03, pDR02, pDR07 and pDR09 |
| rhlI_Fwd_HindIII | CAGTAAGCTTAGAACATCCAGAA GAAGTTCG | PAO1 gDNA | pSR04 |
| rhlI_Rev_BamHI | CAGTGGATCCTCATACTTCCAGC GATTCAGAGAG | PAO1 gDNA | pSR04 |
| rhlR_Fwd_HindIII | TAACGAAAGCTTCCTGCAGGGC GACTTCTAC | PAO1 gDNA | pDR03, pDR05 |
| rhlR_ Rev_BamHI | TTGCTGGATCCTCACTGCATCTG GTATCGCTCC | PAO1 gDNA | pDR03, pDR05 |
| rhlA_Fwd_HindIII | CAGTAAGCTTCCTGGGCAAGAG CACCTACG | PAO1 gDNA | pSR06, pDR04, pDR06 |
| rhlA_Rev_BamHI | CAGTGGATCCTCATATGCAAACC GATACCAACAGA | PAO1 gDNA | pSR06, pDR04, pDR06 |
| rpsL_Fwd_HindIII | CAGTAAGCTTGTACCGGTCTGG CTTACCAC | PAO1 gDNA | pSR07 and pDR08 |
| rpsL_Rev_BamHI | CAGTGGATCCTCAGTGTGCCGA GTTCGGCTTTT | PAO1 gDNA | pSR07 and pDR08 |
| rpoD_Fwd_HindIII | CAGTAAGCTTTCGTCCAGGAGA ACCTTGAA | PAO1 gDNA | pSR08 |
| rpoD_Rev_BamHI | CAGTGGATCCTCATGTCTCGAAT ACGTTGATCC | PAO1 gDNA | pSR08 |

| | | | |
|---|---|---|---|
| recA_Fwd_HindIII | CAGTAAGCTTAGTGAGCGCTGC CAGTTC | PAO1 gDNA | pSR09 |
| recA_Rev_BamHI | CAGTGGATCCTCAGAATTGGCG TTCGATCTGTC | PAO1 gDNA | pSR09 |
| **For cloning promoter region upstream of GFP** | | | |
| lasI_Fwd_Kpn1 | CAGTGGTACCGCCCGGAAGGCC ATGTTTTG | PAO1 gDNA | pDR01 |
| lasI_Rev_Pac1 | CAGTTTAATTAATCATGGGCCAG TGGTATCGAGAAT | PAO1 gDNA | pDR01 |
| lasR_Fwd_Kpn1 | ATGACGGTACCTGGAAAAGTGG CTATGTCGC | PAO1 gDNA | pDR02 and pDR05 |
| lasR_Rev_Pac1 | CGTTCTTAATTAATTAGGCGCTC CACTCCAATTTTC | PAO1 gDNA | pDR02 and pDR05 |
| lasB_Fwd_Kpn1 | CAGTGGTACCCGAATTCCGCGG CCAGGAAAGCGTGCAA | PAO1 gDNA | pDR06 and pDR09 |
| lasB_Rev_Pac1 | CAGTTTAATTAATCAATCGAACA ACAGGTCAAGCGTA | PAO1 gDNA | pDR06 and pDR09 |
| rhlI_Fwd_Kpn1 | CAGTGGTACCAGAACATCCAGA AGAAGTTCG | PAO1 gDNA | pDR03 |
| rhlI_Rev_Pac1 | CAGTTTAATTAATCATACTTCCA GCGATTCAGAGAG | PAO1 gDNA | pDR03 |
| rhlR_Fwd_Kpn1 | TAACGAGGTACCCCTGCAGGGC GACTTCTAC | PAO1 gDNA | pDR04 |
| rhlR_Rev_Pac1 | TTGCTTTAATTAATCACTGCATCT GGTATCGCTCC | PAO1 gDNA | pDR04 |
| rhlA_Fwd_Kpn1 | CAGTGGTACCCCTGGGCAAGAG CACCTACG | PAO1 gDNA | pDR07 |
| rhlA_Rev_Pac1 | CAGTTTAATTAATCATATGCAAA CCGATACCAACAGA | PAO1 gDNA | pDR07 |
| rpsL_Fwd_Kpn1 | CAGTGGTACCGTACCGGTCTGG CTTACCAC | PAO1 gDNA | pDR08 |
| rpsL_Rev_Pac1 | CAGTTTAATTAATCAGTGTGCCG AGTTCGGCTTTT | PAO1 gDNA | pDR08 |
| **Construction of double fluorescent gene reporters** | | | |
| GFP_Fwd_ KpnI_PacI | CAGTGGTACCCCTACCTTAATTA A GCCGCTTTAAGAAGGAGGTA | pEX-A128-GFP | pUC18-mini-Tn7-Gm-mCherry-GFP |
| GFP_Rev_ HindIII_SpeI | CAGTAAGCTTGGGATGACTAGTT CATAACCCCGCTACTCATCATTT G | pEX-A128-GFP | pUC18-mini-Tn7-Gm-mCherry-GFP |
| Fwd_Term_Nsi1 | CAGTATGCATTAATAATAAACGC AGAAAGGC | pEX-A128-terminators | pUC18-mini-Tn7-Gm-mCherry-GFP |

| | | | |
|---|---|---|---|
| Rev_Term_Sac1 | CAGTGAGCTCTGCAGGTCGTCT CGGATC | pEX-A128-terminators | pUC18-mini-Tn7-Gm-mCherry-GFP |
| **Colony PCR** | | | |
| Tn7L_rev[a] | GGGTGTAGCGTCGTAAGCTAAT | *E. coli* colonies | Colony PCR to check insertion in *E. coli* |
| mCherry_rev2[b] | GGATATCCGCTGGGTGTTTA | *E. coli* colonies | Colony PCR to check insertion in *E. coli* |
| PTn7L | ATTAGCTTACGACGCTACACCC | *P. aeruginosa* colonies | Colony PCR to check insertion in *P. aeruginosa* |
| PglmS-up | CTGTGCGACTGCTGGAGCTGA | *P. aeruginosa* colonies | Colony PCR to check insertion in *P. aeruginosa* |

[a]To confirm insertion of promoter region upstream of GFP in the mini-Tn7 vector, colony PCR was performed using Tn7L_rev primer, together with the corresponding “Rev_PacI” promoter-specific primer.

[b]To confirm insertion of promoter region upstream of mCherry in the mini-Tn7 vector, colony PCR was performed using mCherry_rev2 primer, together with the corresponding “Fwd_HindIII” promoter-specific primer.
